# Fluid forces control structural remodeling of blind-ended lymphatic microvessels

**DOI:** 10.1101/2025.04.14.648799

**Authors:** Jacob C. Holter, Shashwat S. Agarwal, Joseph W. Tinapple, Joseph M. Barlage, Travis H. Jones, Jonathan W. Song

## Abstract

The transport function of lymphatic vessels is altered during tissue injury, inflammation, and cancer. Defects in lymphatic function are associated with changes to the biophysical microenvironment, including pressure and flow. However, the ability of fluid forces to orchestrate the remodeling of blind-ended lymphatic vessels and lymphangiogenesis is not well understood. Here, we developed a novel microphysiological system (MPS) that recapitulates the blind-ended microanatomy and fluid absorption properties of capillary lymphatics. Our MPS implements a continuum of pressure-driven interstitial, transmural, and luminal flow to mimic fluid forces naturally present within the lymphatic microenvironment. We found that interstitial flow (IF) and VEGF-C cooperate during lymphangiogenesis. Notably, we observed that sprouting was most prominent at the blind-ended region of lymphatic vessels where transmural flow was highest in our MPS. Moreover, IF guided invading sprouts into the surrounding ECM antiparallel to streamlines within a non-uniform 3-D flow field. Strikingly, flow-induced elongation and axial alignment of intraluminal cells propagated to vessel-level phenotypic differences, such as vasoconstriction and helical patterning. The structural remodeling of these lymphatic vessels was concurrent with lymphangiogenesis. Thus, our results reveal how extravascular and intraluminal endothelial cells integrate signals from native fluid forces to coordinate the expansion and remodeling of capillary lymphatics.

## 1. Introduction

The lymphatic vascular system is comprised of a network of one-way conduits that collect and transport interstitial fluid containing proteins, lipids, and cells necessary for tissue fluid homeostasis and immune surveillance.^[1]^ Initial lymphatics, or capillary lymphatics, are the most distal blind-ended vessels of the lymphatic system where fluid uptake begins. Interstitial, transmural, and luminal flow are fluid mechanical forces naturally present within the microenvironment of capillary lymphatics.^[2]^ Interstitial fluid pressure (IFP) drives interstitial flow (IF) within tissue spaces, and fluid clearance is initiated as transmural flow crosses the lymphatic endothelium. This drained fluid is then transported as luminal flow into the pre-collecting and collecting lymphatics and eventually propelled back to blood circulation.^[3]^ In addition to conferring mechanical stimuli, IF can help shape local gradients of growth factors and other biomolecules within the extracellular matrix (ECM).^[4]^ Elevated IF originating from leaky blood vessels is associated with a number of pathological conditions including inflammation and cancer, and also occurs during wound healing.^[5]^ Consequently, lymphatic expansion and lymphangiogenesis are characteristic of dermal tissue regeneration,^[6]^ chronic inflammation,^[7]^ and metastasis^[8]^ in many human and murine tumors where elevated IFP and IF are present. Further, transmural and luminal flow have been shown to be important regulators of lymphatic endothelial function.^[9]^

The rise of *in vitro* microphysiological systems (MPS), or lymphatic-on-chip platforms, has made it possible to replicate tissue microenvironments with tractable control over experimental parameters and systematically study lymphatic biology.^[10]^ Using MPS, the regulatory role of IF in promoting microvascular networks^[11]^ and sprouting morphogenesis has been demonstrated for blood^[12]^ and lymphatic endothelial cells (LECs).^[13]^ Moreover, MPS have been leveraged to reveal how IF regulates lymphatic junction morphogenesis similar to the button-like junctions of capillary lymphatics.^[14]^ Other *in vitro* systems have successfully cultured three-dimensional (3-D) human lymphatic vessels and integrated various stromal components such as high-density collagen,^[15]^ breast cancer-associated fibroblasts (CAFs),^[16]^ breast cancer cells,^[17]^ and head and neck cancer (HNC) spheroids^[18]^ to model the tumor microenvironment (TME). And lymphatic dysfunction has been recapitulated for acute and chronic inflammation.^[19]^ Nevertheless, these systems typically feature open-ended lumen structures that do not recreate the blind-ended microanatomy of capillary lymphatics and resultant flow dynamics. Furthermore, IF is often administered unidirectionally towards two-dimensional (2-D) cell-ECM interfaces at discrete levels of velocity (e.g., 1 or 3 μm/s),^[20]^ which does not recreate the range of flow regimes exerted on a single *in vivo* lymphatic microvessel. Among the few studies that have reconstructed blind-ended lymphatics, investigation centered on drainage functionality for tissue scaffolds^[21]^ and breast cancer cell escape.^[22]^

While the recapitulation of native fluid forces has been a focus of lymphatic-on-chip platforms, to date, no system has integrated 3-D lumen structures,^[22-23]^ blind-ended microanatomy,^[24]^ and physiological IF^[5b]^ to advance our understanding of lymphatic biology and pathophysiology. Using a xurography-based microfabrication technique previously developed by our group,^[25]^ we engineered a novel lymphatic-on-chip platform that recapitulates a lymphatic capillary and its native biophysical microenvironment. By implementing physiological interstitial fluid flow, we generated continuous transverse, or transmural, flow across the vessel wall to mimic lymphatic fluid transport and model lymphangiogenesis. In doing so, we demonstrated the stimulatory effects of IF and vascular endothelial growth factor C (VEGF-C) on sprouting morphogenesis in a 3-D blind-ended vessel. Thus, we highlight the synergistic capacity of IF and VEGF-C on lymphangiogenesis when applied together and the relative contributions of each when decoupled. Convective IF was sufficient to induce and sustain sprouting independent of VEGF-C. Furthermore, transmural flow velocity significantly correlated with lymphangiogenic activity, which suggests that transmural flow is a major inductive cue for sprouting. Beyond lymphangiogenesis, we investigated the morphological changes of intraluminal cells due to flow. Within a 3-D microtissue context, we propose that cell-level responses to interstitial, transmural, and luminal flow manifest at the vessel level insofar as vessels remodel by constriction and helical patterning. These results showcase a coordinated structural remodeling of capillary lymphatics and suggest that there is a profound interrelationship between intra-and extraluminal regions—mediated by flow dynamics. Given the role of tissue geometry on branching morphogenesis,^[26]^ our MPS expands upon these insights and can further reveal novel lymphangiogenic mechanisms that govern lymphatic transport and function.

## 2. Results

### 2.1. Engineering a blind-ended lymphatic microvessel *in vitro*

To create blind-ended human lymphatic vessels, we first designed a novel microfluidic device constructed from a layer-by-layer assembly of poly(dimethylsiloxane) (PDMS) (**Figure S1**, Supporting Information) and xurography techniques established by our lab.^[25]^ Briefly, a cylindrical lumen was templated by placing a nitinol wire within a 3-D hydrogel chamber such that the wire was elevated above the base, confined within a microchannel, and extended to the midplane of the chamber. Subsequently, 3 mg mL^-1^ collagen was pipetted into the gel port and polymerized around the wire to cast a hollow lumen structure upon wire removal. Human dermal lymphatic endothelial cells (HDLECs) were then seeded into the lumen to form a blind-ended lymphatic microvessel (**Figure 1A-B**). The rigidity of the 250-µm-diameter wire, coupled with the layer-by-layer design, permitted the lumen to be cast on a specified *z*-plane within the MPS to facilitate imaging and—in later iterations—ensure symmetrical flow dynamics. Moreover, the cylindrical shape of the wire enabled the lumen to be templated with a circular profile providing more physiologically relevant geometry and wall shear stresses^[27]^ than rectangular microvessel analogues.^[28]^ The Static device (**Figure 1C**) consisted of 4-mm-diameter inlet and outlet ports, two 1.5-mm-diameter gel ports, and a 4-mm-diameter central gel chamber.

**Figure 1.**
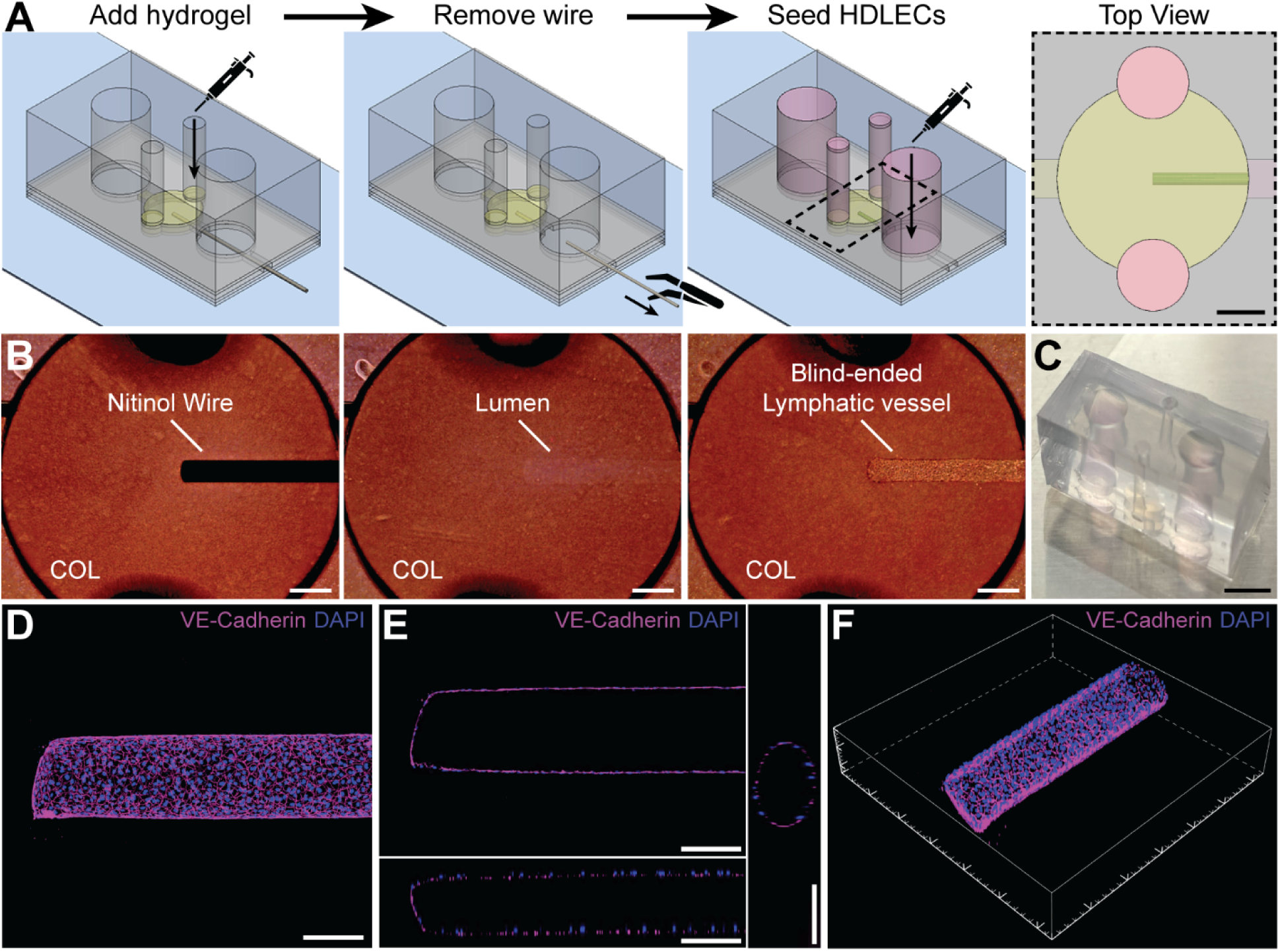
Engineered blind-ended lymphatic microvessel. A) Schematic of the novel PDMS-based Static microdevice and subsequent steps to generate a lymphatic vessel. A 250-μm-diameter wire is placed within the device terminating at the midplane of the central ECM chamber. Collagen-based hydrogel is pipetted into the central chamber via the lateral gel ports. Upon gel polymerization, the wire is removed to expose a blind-ended hollow lumen structure. The templated lumen is then seeded with human dermal lymphatic endothelial cells (HDLECs) to create a microvessel. PDMS layers are represented as transparent gray, glass slide as blue, collagen as yellow, cell culture media as pink, and HDLECs as green. Black dashed region depicts the top view of the central gel chamber with the engineered microtissue fully constituted. Scale bar is 1 mm. B) Phase-contrast images corresponding to steps in A to illustrate workflow resulting in a representative blind-ended lymphatic vessel encased in 3 mg mL^-1^ collagen (COL). Scale bars are 500 μm. C) Photograph of Static microfluidic device. Scale bar is 5 mm. D) Confocal *z*-projection of a representative blind-ended lymphatic microvessel stained for junction protein VE-Cadherin (antibody, magenta) and nuclei (DAPI, blue). E) Orthogonal views to demonstrate a patent, perfusable microvessel with a monolayer of HDLECs: *x-y*, *x-z*, and *y-z* cross sections (counterclockwise from upper left). Scale bars in D and E are 200 μm. F) 3-D render of a blind-ended lymphatic vessel from confocal *z*-stack. Major tick marks are 200 μm.

We validated the formation of a blind-ended microvessel with a monolayer of HDLECs, consistent with capillary lymphatics,^[5a, 24]^ by employing immunofluorescence and confocal microscopy. The endothelial-specific adherens junction protein, VE-cadherin, was stained to confirm cell-cell contacts, and nuclei were counterstained with DAPI (**Figure 1D**). An intact, confluent monolayer of endothelium with intercellular junctions was observed, thereby confirming microvessel integrity. Orthogonal views and a 3-D render of a representative lymphatic capillary illustrate the blind-ended geometry and elliptical cross section engineered into the patent microvessel (**Figure 1E-F**). These lymphatic vessels were cultured under static, no flow conditions for four days.

### 2.2. Reconstituting the biophysical microenvironment of a functional lymphatic capillary

Next, we recapitulated the native biophysical microenvironment that regulates interstitial and lymphatic fluid transport. This environment includes the movement or convection of interstitial fluid due to flow-driving pressure gradients, thereby enabling continuous transverse fluid absorption by blind-ended lymphatic capillaries.^[29]^ Fluid pressure gradients and ECM hydraulic resistance determine IF levels in native tissue.^[5b]^ To this end, we modified the Static device (**Figure 1**) into a Flow device variant (**Figure 2**) to independently control these parameters and apply defined levels of IF. The source of upstream fluid pressure was afforded by a liquid hopper, which was attached to the inlet of the device and filled with cell culture media to a specified height of hydrostatic pressure. The liquid hopper was constructed from a sterile 20-mL syringe whose tip interfaced with the 4-mm-diameter inlet by a simple press fit connection. We then modulated the length of the upstream microchannel of the device and the resultant pressure gradient (**Figure 2A**). The collagen-filled microchannel provided the principal resistive element of the microdevice due to its dimensions^[30]^ and the hydraulic resistance of the collagen gel, defined by Darcy’s law.^[5b]^ Given the substantial resistance of the microchannel relative to the ECM chamber and vessel, the drop in IFP occurred primarily within the upstream channel as a means of controlling IF (**Figure S2A**, Supporting Information). Lastly, a ramp was cut at the outlet of the PDMS microdevice with a sterile blade to allow cell culture media to flow out the back of the device to zero gauge pressure.^[31]^ Thus, a continuous liquid circuit was formed by a liquid ramp connecting the media-filled outlet to pooled media on the surface of the glass slide— a hydraulic analogue to electrical grounding (**Figure 2B**).

**Figure 2.**
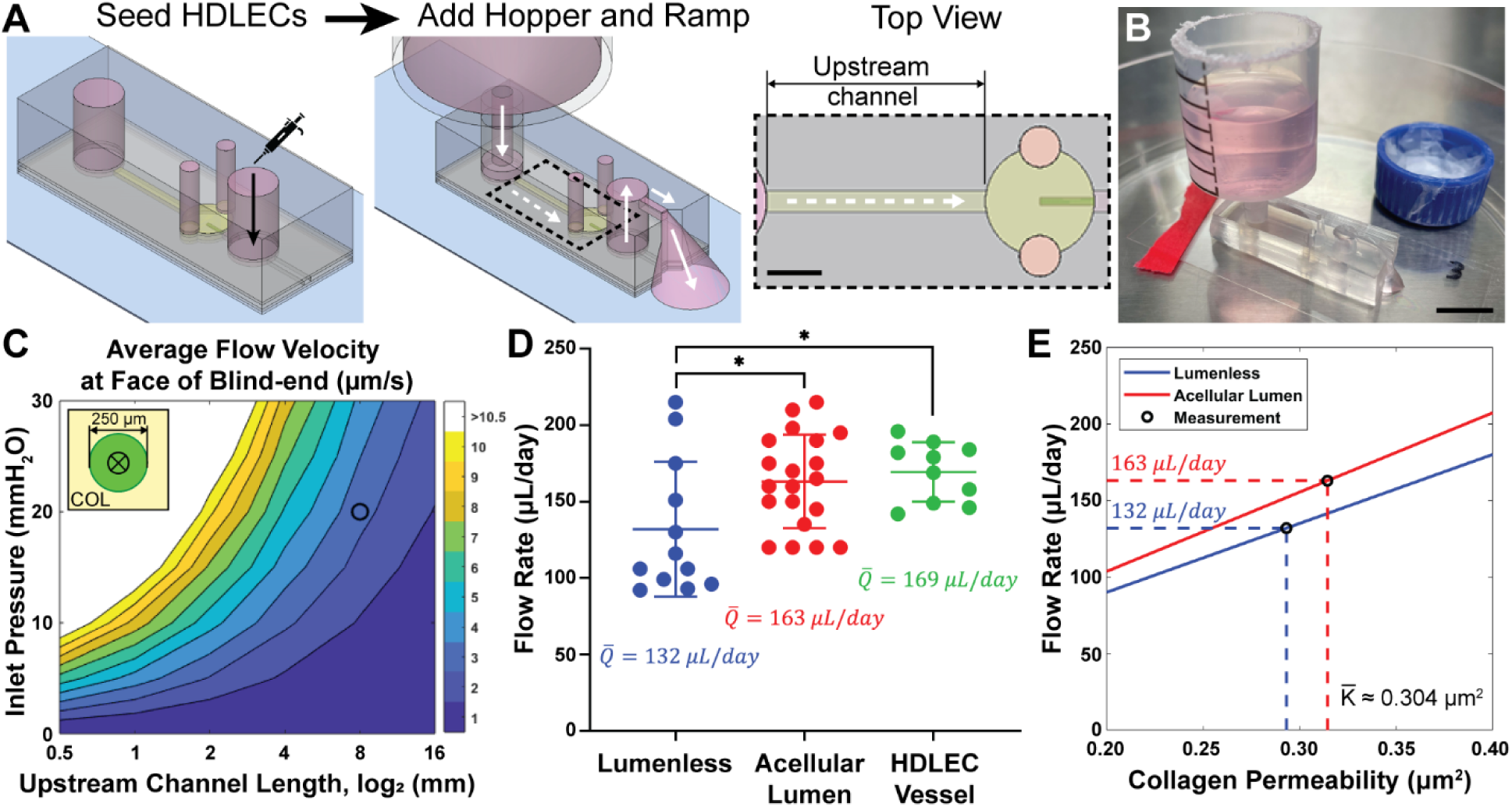
Mimicking the biophysical microenvironment of a functional lymphatic capillary. A) The Flow device was engineered to accommodate interstitial flow (IF) by increasing the length of the upstream microchannel for resistance and integrating a liquid hopper for pressure. After seeding HDLECs, a liquid hopper was inserted into the inlet port and filled with cell culture media to a specified height. A ramp was cut at the outlet port to allow perfused media to flow out the back of the device to zero gauge pressure, analogous to electrical grounding. White arrows depict the direction of media flow and dashed white arrows represent IF. Black dashed region displays the top view of the MPS with modular upstream channel length. Scale bar is 2 mm. B) Photograph of a fully assembled Flow device contained within a Petri dish to avoid contamination, improve humidity, and minimize evaporation. Inverted blue cap is filled with 1× PBS to help maintain humidity. C) Finite element analysis (FEA) results for average flow velocity at the face of the blind-ended lymphatic vessel when varying upstream channel length and inlet pressure head. The black circle denotes the selection of an 8 mm channel and 20 mmH_2_O IFP to achieve a flow velocity of ∼ 3 μm/s. Inset is a schematic of the circular blind-ended face of the 250-μm-diameter vessel surrounded by collagen (COL) where ® denotes fluid flow into the vessel. D) Empirical daily flow rates (Q-) by measuring pass-through volume of cell culture media for three different configurations of the Flow device to validate FEA results. Data are expressed as mean ± SD. One-way ANOVA was performed with Tukey pairwise comparisons, where * indicates *p*-value < 0.05. E) Estimation of the bulk hydraulic permeability (K-) of semi-porous 3 mg mL^-1^ collagen hydrogel given the measured flow rates and FEA results.

The placement of the liquid hopper directly opposite the lymphatic vessel generated impinging IF that then became transmural flow with highest velocity at the blind-ended region. To calculate the average theoretical velocity at the face of the vessel, we employed finite element analysis (FEA) (**Figure 2C**). Among the various arrangements, we proceeded with an 8-mm-long upstream channel and 20 mmH_2_O of inlet pressure to yield an average transverse velocity across the blind-ended surface of 3 μm/s—representative of elevated *in vivo* IF velocity.^[4, 20a]^ We then experimentally determined the hydraulic permeability of collagen by measuring flow rates (Q-) of media through three different configurations of the Flow device: i) lumenless, ii) acellular lumen, and iii) HDLEC vessel (**Figure 2D**). Flow rate experiments used the same upstream channel length and inlet pressure as our selected FEA model: 8 mm and 20 mmH_2_O, respectively. The lumenless Flow device was constructed without templating a lumen, and hence, lacked cylindrical void space within the collagen. The average pass-through flow rates for the lumenless, acellular lumen, and HDLEC vessel configurations were 132, 163, and 169 μL/day, respectively. As expected, the lumenless Flow device exhibited the largest hydraulic resistance of the three configurations, evident by the lowest flow rate. Moreover, similar flow rates for the acellular lumen and HDLEC vessel configurations confirmed that the collagen-filled upstream channel was the dominant resistive element within the MPS. Experimental flow rates for the lumenless and acellular lumen cases were then compared to FEA results to determine the hydraulic permeability of 3 mg mL^-1^ collagen (**Figure 2E**). The two configurations independently converged on an average permeability (K-) of 0.304 μm^2^, which is consistent with previously reported measurements from literature^[5b, 32]^ and our lab.^[33]^ This empirically estimated collagen permeability was used as an input for ensuing computational modeling.

### 2.3. Interstitial flow initiates and sustains lymphangiogenesis independent of VEGF-C

To identify the effects of the native biophysical microenvironment on lymphangiogenesis, we compared lymphatic sprouting for four different conditions: (i) Static, (ii) Flow, (iii) Static+VEGF-C, and (iv) Flow+VEGF-C (**Figure 3A**). For the two growth factor conditions, 50 ng mL^-1^ of exogenous VEGF-C was added to cell culture media and administered to the inlet, outlet, and liquid hopper. Thus, IF facilitated mass transport of VEGF-C by convection in the Flow+VEGF-C condition, and diffusion enabled VEGF-C transport in the Static+VEGF-C condition. Treatment with IF and/or VEGF-C commenced at Day 1 to allow LECs to adhere to the lumen surface and stabilize as a monolayer over the first 24 hours after seeding on Day 0. All vessels were then cultured to Day 4 before subsequent fixation and immunofluorescent staining (**Figure S3**, Supporting Information). For the Static condition, we observed virtually no lymphatic sprouts: 0.2 ± 0.1% sprouting area, as a percentage of vessel surface area. In contrast, the Flow condition prompted significant sprouting against the direction of IF compared to Static: 10.5 ± 3.1% (**Figure 3B**). Static+VEGF-C induced a sprouting area of 1.1 ± 0.7%, differentiating it from the Static condition, although 10-fold less than Flow. Flow+VEGF-C promoted copious lymphangiogenesis, resulting in a sprouting area of 30.0 ± 9.2% or 3-fold more than Flow alone. These results confirm previous reports that VEGF-C augments lymphangiogenesis in the presence of IF.^[20b]^

**Figure 3.**
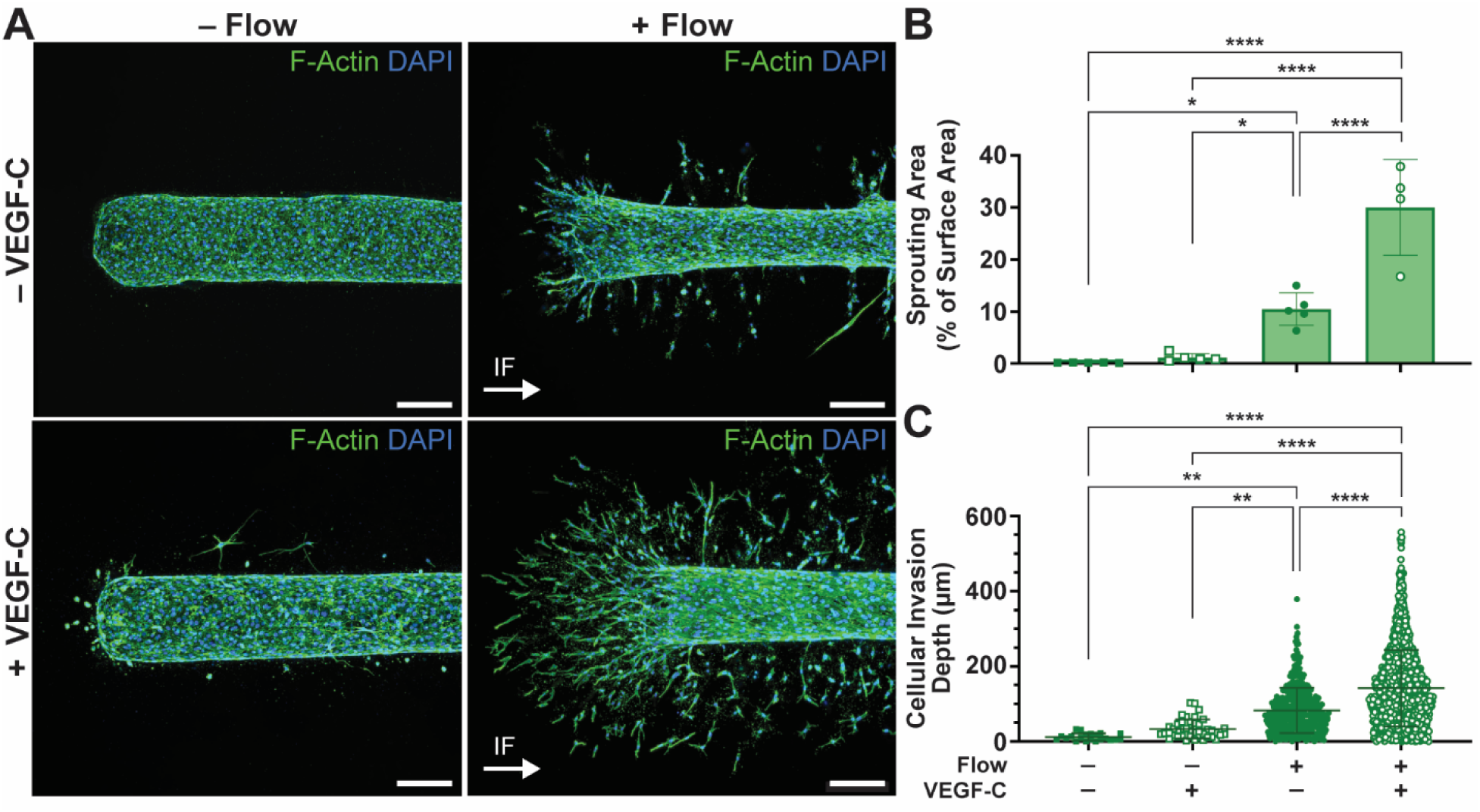
Interstitial flow initiates and sustains lymphangiogenesis independent of VEGF-C. A) Matrix of representative lymphatic microvessels subjected to the four experimental conditions and fixed at Day 4. Images are confocal *z*-projections of vessels stained for F-actin (phalloidin, green) and nuclei (DAPI, blue). White arrows indicate the direction of physiologically relevant interstitial flow (IF). Scale bars are 200 μm. B) Quantification of sprouting area per vessel as a percentage of total vessel surface area (*N* = 4 or 5 vessels). C) Invasion depth of individual sprouting LECs into the surrounding ECM measured from the vessel surface (*N* > 20 cells for **–** Flow conditions, and *N* > 400 cells for +Flow conditions). Data are expressed as mean ± SD. One-way ANOVA was performed with Tukey pairwise comparisons, where * indicates *p*-value < 0.05, ** < 0.01, *** < 0.001, and **** < 0.0001.

In addition to sprouting area, we evaluated the invasion depth of individual sprouting cells into the ECM across the four experimental conditions (**Figure 3C**). As anticipated, invasion depth results followed a similar pattern to sprouting area. For the Static condition, many vessels had no LEC invasion, while the few cells that did invade averaged only 12.0 ± 9.8 μm. In contrast, Flow LECs invaded an average of 82.8 ± 60.0 μm, significantly greater than those under Static conditions. Static+VEGF-C vessels consistently exhibited some number of LECs to invade a shallow depth of 33.5 ± 25.1 μm, distinguishing it from Static-only treatment, but significantly less than Flow. The most significant invasion occurred for Flow+VEGF-C vessels: 142.2 ± 101.6 μm. Collectively, by quantifying sprouting area and invasion depth across the four conditions, we demonstrated that interstitial flow initiates and sustains lymphangiogenesis into the perivascular ECM independent of VEGF-C.

### 2.4. Zonal sprouting analysis reveals selective lymphangiogenesis at the blind end

We next evaluated regional differences in sprouting given the heightened IF velocity at the blind end of our lymphatic microvessels. 2-D projections from confocal *z*-stacks of the vessels were divided into five zones: Zone 0 representing the 250-μm-diameter blind-ended front face of the vessel, and Zones 1–4 corresponding to 250-μm-wide segments along the length of the vessel axis. Zones were taken to be the extraluminal ECM space normal to the microvessel walls (**Figure 4A**). Transverse flow velocity, intraluminal shear stress, and normalized sprouting area were then analyzed for all five zones at each experimental condition.

**Figure 4.**
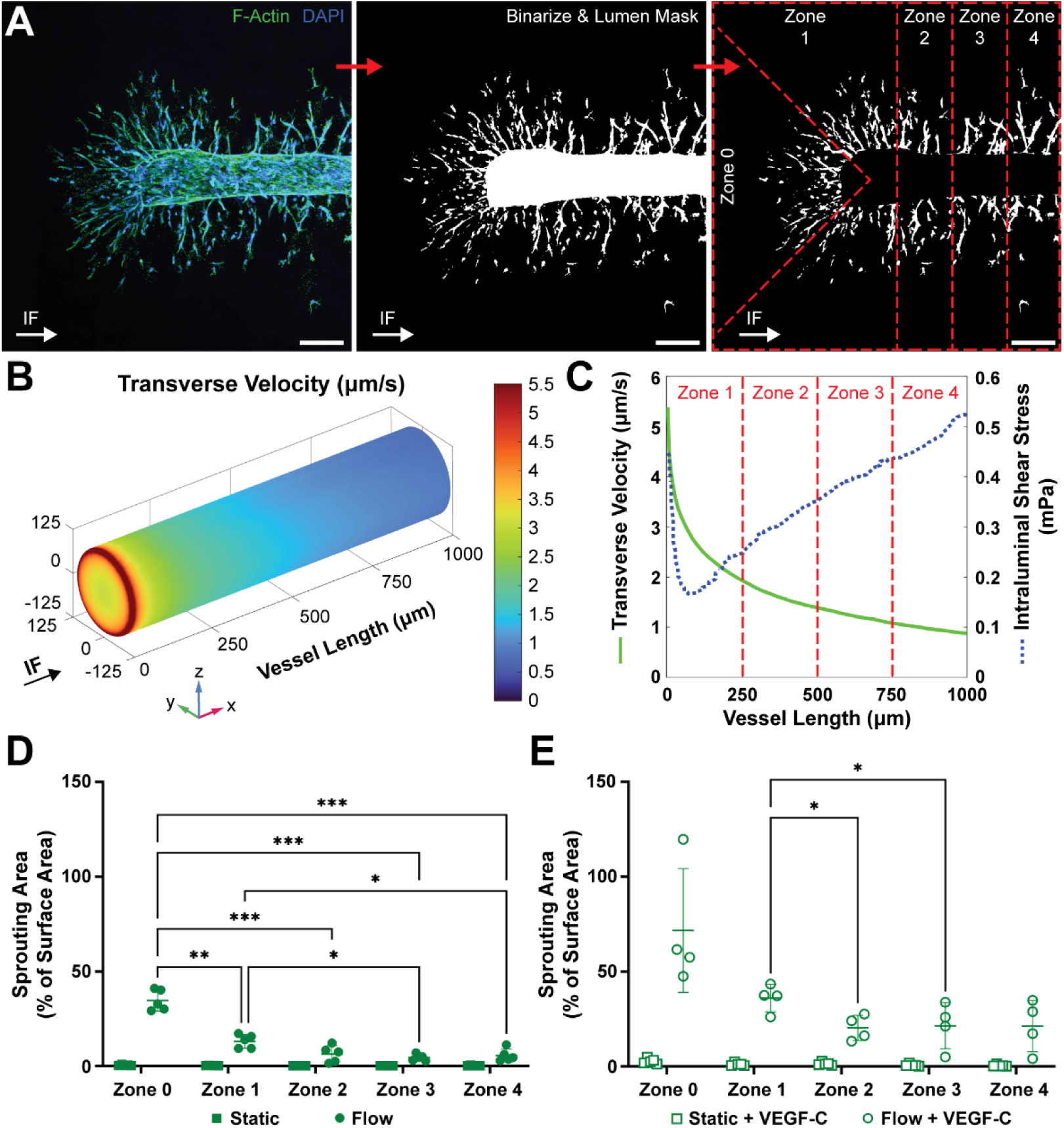
Selective lymphangiogenesis occurs at the blind-end due to elevated local transverse velocity. A) Confocal *z*-projection images of vessels were binarized, and sprouts were isolated. A map divides a representative vessel into five zones (Zones 0–4) depicted by red dashed lines corresponding to 250-μm vessel segments normal to the vessel surface. Scale bars are 200 μm. B) Heat map of transverse flow velocity across the lymphatic vessel surface from finite element analysis (FEA). Velocity is greatest at the blind end due to the orientation of the vessel pointing towards the source of flow. C) Transverse velocity and intraluminal shear stress from FEA plotted along the length of the vessel, given axial symmetry. Demarcated zones highlight varying flow velocity and shear stress for different regions of the same microvessel due to the blind end. D and E) Sprouting area per zone normalized as a percentage of respective vessel surface area within each zone, with and without VEGF-C (*N* = 4 or 5 vessels). Sprouting area— as a metric for lymphangiogenic activity—strongly correlates with transvascular flow velocity and negatively correlates with intraluminal shear stress. Data are expressed as mean ± standard deviation. Two-way ANOVA was performed with Tukey pairwise comparisons, where * indicates *p*-value < 0.05, ** < 0.01, and *** < 0.001.

Firstly, using FEA (**Figure 2C**), we confirmed that transmural flow velocity was highest at the blind end due to its proximity to the upstream channel and the resultant impinging flow (**Figure 4B**). A peak velocity of 5.5 μm/s was observed at the edges of the blind-ended face. Transmural velocity then diminished moving down the length of the vessel, ultimately reaching ∼ 1 μm/s at a distance one millimeter from the front face (**Figure 4C**). Within each zone, a range of transverse flow velocities across the vessel wall was numerically solved: Zone 0: 5.5–3.1 μm/s; Zone 1: 5.5–1.8 μm/s; Zone 2: 2.0–1.2 μm/s; Zone 3: 1.4–1.0 μm/s; and Zone 4: 1.1–0.8 μm/s. Comparatively, intraluminal shear stress was lowest in Zone 1, towards the blind end where intravascular flow begins, and increased moving down the vessel. Non-uniform wall shear stress is consistent with additional FEA results that show intraluminal flow to accelerate downstream while moving away from the blind end (**Figure S2B**, Supporting Information).

For both Flow and Flow+VEGF-C conditions, the largest amount of zonal sprouting occurred at Zone 0: 34.2 ± 5.7% and 71.6 ± 32.6%, respectively (**Figure 4D-E**). Sprouting then diminished for each subsequent zone moving down the vessel axis away from the source of IF. For the Flow condition, Zone 0 had statistically more sprouting than any of the other four zones, while Zone 1 had significantly more sprouting than downstream Zones 3 and 4. For Flow+VEGF-C treatment, more sprouting occurred at Zone 0 than any of the other four zones, while Zone 1 had significantly more sprouting than downstream Zones 2 and 3. Observed sprouting area per zone was then correlated to the previously determined transverse flow velocity and shear stress. Significant Pearson correlation coefficients of 0.937 and 0.940 were quantified for average transverse flow per zone (Zones 0–4) to normalized sprouting area, for Flow and Flow+VEGF-C conditions, respectively (*p*-values of 0.019 and 0.017, respectively). Comparatively, Pearson correlation coefficients of -0.815 and -0.735 were quantified for average shear stress per zone (Zones 1–4) to normalized sprouting area, for Flow and Flow+VEGF-C conditions, respectively (*p*-values of 0.185 and 0.265, respectively). Thus, we show that selective lymphangiogenesis occurred at the blind end of the microvessel positively correlating with transvascular flow and negatively correlating with shear stress.

Furthermore, we evaluated the zonal distribution of lymphangiogenesis by assessing sprouting area per zone as a percentage of total sprouting area for each vessel. While the addition of VEGF-C led to increased sprouting for each respective zone when comparing IF conditions (**Figure 4D-E**), the spatial distribution of sprouts between Flow and Flow+VEGF-C was not statistically significant for each of the five zones (**Figure S4**, Supporting Information). Thus, VEGF-C augmented total flow-potentiated lymphangiogenesis, however, the zonal distribution of sprouting remained the same in the presence or absence of VEGF-C.

### 2.5. Lymphangiogenic sprouts are guided by interstitial flow streamlines

The ability of IF to guide the direction of invading lymphatic sprouts was assessed. Firstly, IF velocity and directional streamlines within the collagen gel were determined by FEA (**Figure 5A**). Examination of the collagen matrix at the *x-y* midplane of the vessel confirmed concentrated flow and augmented velocity at the blind end, reaching a peak magnitude of 5.5 μm/s. Given the asymmetrical orientation of the vessel pointing towards the source of fluid flow, the highest IF velocities occurred in the gel region near the blind end and diminished further downstream. Streamlines revealed IF to move towards and ultimately enter the conductive vessel normal to its surface—independent of inlet pressure modifications. Utilizing Cellpose,^[34]^ we constructed masks of individual cells that had invaded into the surrounding collagen-based ECM. The major axis of each cell was then compared to the numerically obtained streamline direction of IF at the centroid location of the cell (**Figure 5B**).

**Figure 5.**
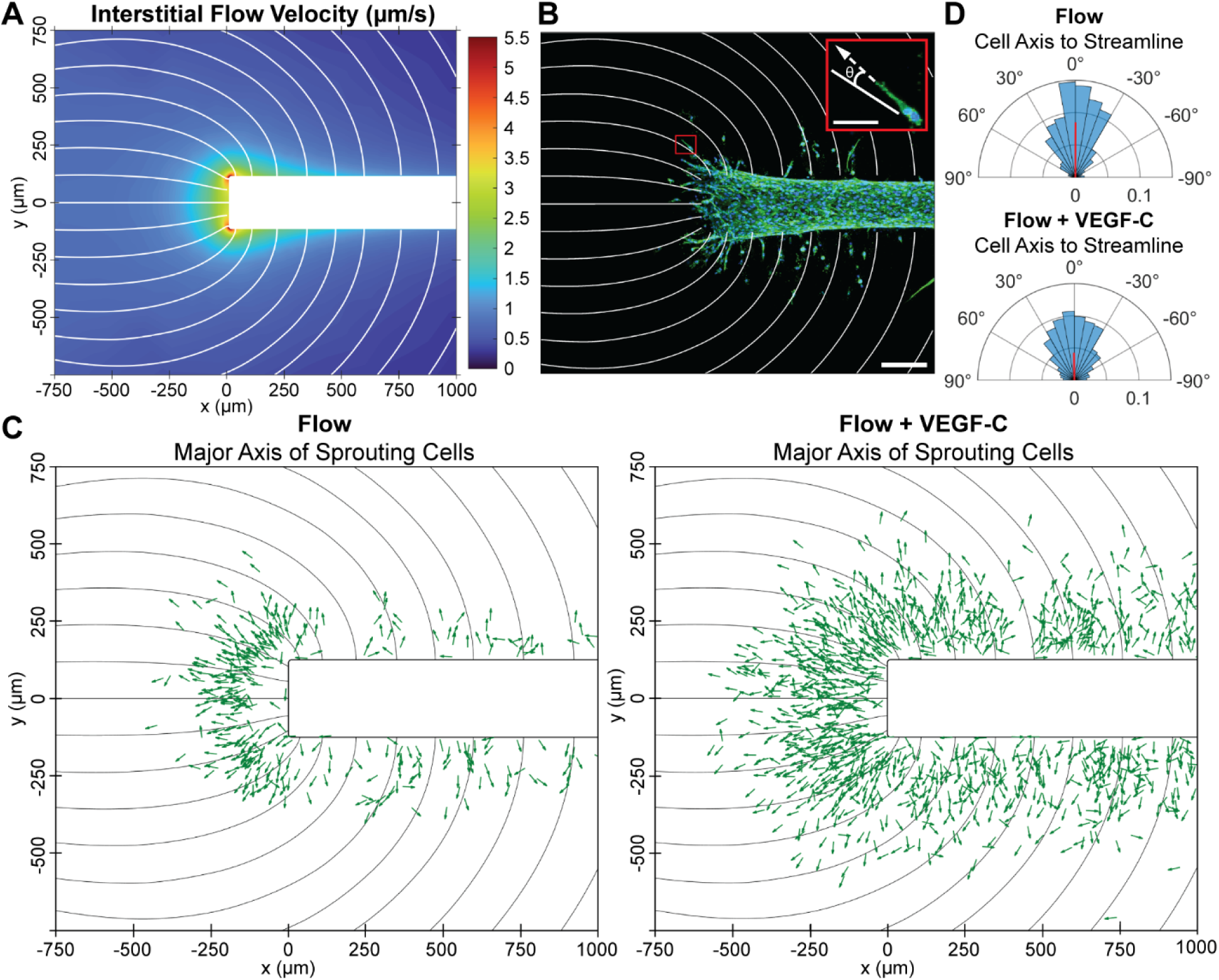
Lymphangiogenic sprouts are guided by interstitial flow (IF) streamlines. A) Magnitude of IF velocity and directional streamlines (white lines) within the collagen gel are plotted for a theoretical lymphatic vessel in our Flow device by finite element analysis (FEA). B) Confocal *z*-projection of a representative microvessel under the Flow condition with FEA streamlines superimposed. Scale bar is 200 μm. Inset is an individual sprouting cell, depicting the angular difference (0_sprout_) between the major axis of the cell (dashed white arrow) and the streamline at the centroid location of the cell (white line). Inset scale bar is 50 μm. C) For Flow and Flow+VEGF-C conditions, the major axes of individual sprouting cells are plotted as green arrows and mapped to their respective location within the collagen matrix (*N* = 4 vessels per condition). Streamlines are displayed as black lines. D) Probability-normalized polar histograms of 0_sprout_ for individual cells in panel C, where 0° is defined as perfectly aligned to the streamline (*N* > 300 cells for Flow, and *N* > 1000 cells for Flow+VEGF-C). Mean resultant vectors denoted as red lines. On average, sprouting LECs invade the ECM collinearly with streamlines, antiparallel to the direction of IF.

The major axis angles of sprouting LECs were mapped to their specific locations within the collagen gel to visualize individual cell orientations relative to streamlines for both Flow and Flow+VEGF-C conditions (**Figure 5C**). The difference between the major axis and the streamline (0_sprout_) was quantified for all invading lymphatic cells and reported as probability-normalized polar histograms (**Figure 5D**). Overall, 71% and 57% of invading LECs were oriented within ± 30° of the streamlines for Flow and Flow+VEGF-C, respectively. Mean angles of 0.21 and 0.73 degrees were calculated for Flow (*N* > 300 cells) and Flow+VEGF-C (*N* > 1000 cells) conditions, respectively. Thus, invading cells oriented themselves—on average— antiparallel to the direction of IF. The mean resultant vector length for Flow was approximately 2-fold greater than that of Flow+VEGF-C, suggesting that IF guides sprouting through the ECM more so than when in conjunction with VEGF-C. We suspect this may be due to the branching of sprouts induced by VEGF-C. In contrast, sprouting cells under the Static+VEGF-C condition were oriented randomly in the absence of flow (**Figure S5**, Supporting Information).

### 2.6. Vessel constriction due to impinging interstitial flow and associated cellular responses

Next, we evaluated changes in lymphatic vessel diameter due to stimulation with IF and VEGF-C. Despite using 250-μm-diameter wires, templated lumens measured approximately 300 μm on Day 0 at steady state. Lumens were then seeded with HDLECs at Day 0, and IF and/or VEGF-C treatment commenced 24 hours later, at Day 1 (**Figure 6A**, **Figure S3A**). All endothelialized vessels constricted ∼ 8% to an average diameter of 276 μm by Day 1 prior to stratified treatment. By Day 4, Static vessels further constricted a total of 17% to 252 μm, primarily occurring over the first 48 hours from seeding. Static+VEGF-C and Flow+VEGF-C vessels had similar daily progression as Static vessels, finishing the experiment at Day 4 with average diameters of 249 μm and 244 μm, respectively. In contrast, by Day 4, Flow vessels significantly constricted a total of 29% to an average diameter of 214 μm, with constriction largely occurring within the first day of IF implementation. Thus, VEGF-C did not have a significant effect on vessel diameter in the absence of IF, while VEGF-C induced vessel dilation in the presence of IF. Though vasodilation is a known property of VEGF-C,^[35]^ we observed that IF was required to mediate changes in vessel diameter.

**Figure 6.**
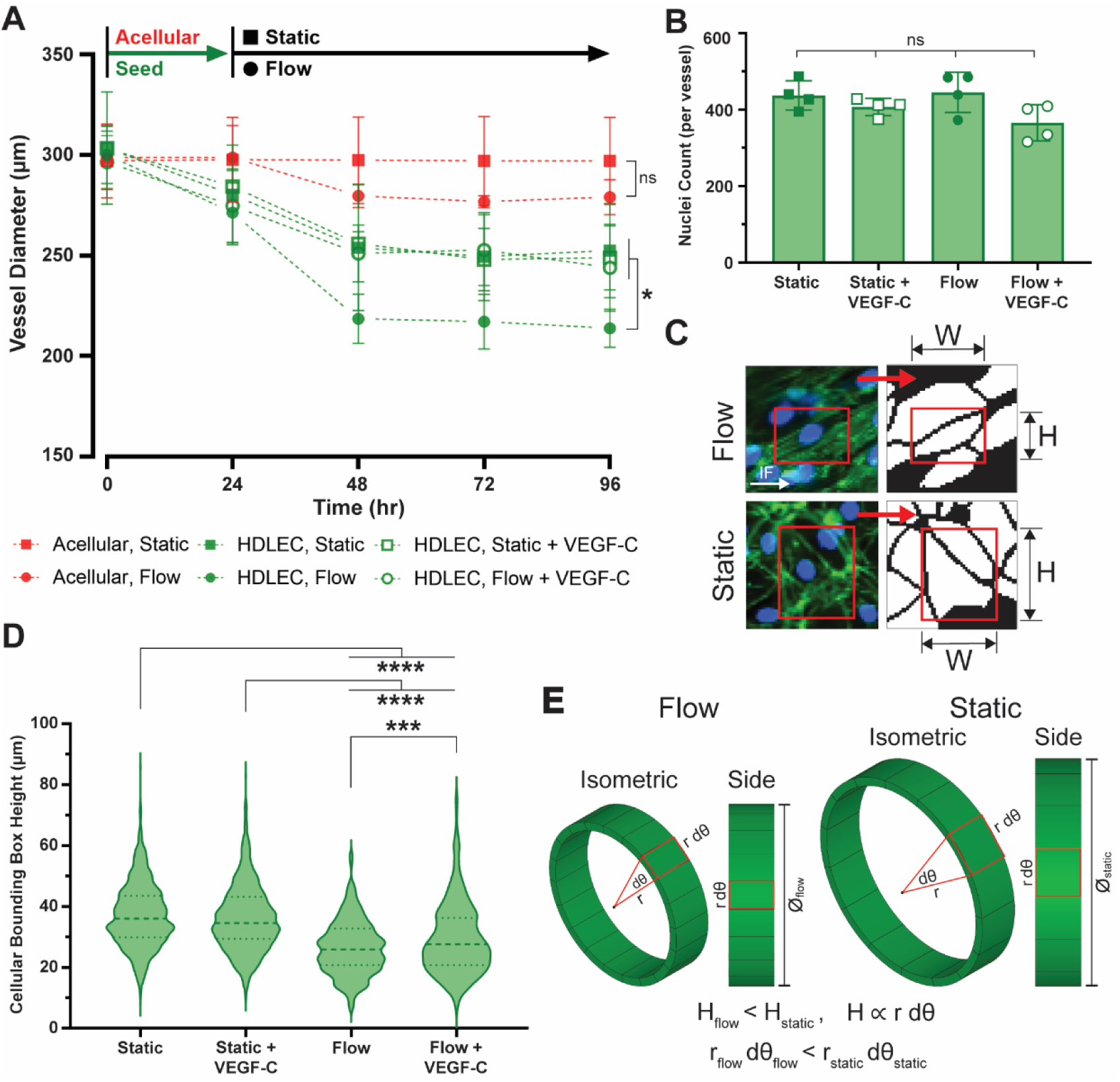
Vessel constriction due to interstitial flow (IF) and associated intraluminal LEC morphology. A) Daily progression of acellular and endothelialized lumens over the course of four days. Lumens are seeded with HDLECs at Day 0 and stimulated with IF and/or VEGF-C at Day 1. Treatment continued for three additional days until the 96 hour mark. In the absence of IF, cells independently constrict lumen diameter, while IF induces further vessel constriction (*N* = 3 lumens for acellular conditions, and *N* ≥ 4 vessels for HDLEC conditions). B) No significant changes in total cell count per vessel were observed across the four vessel conditions (*N* = 4 vessels per condition). C) Representative binary cell segmentation masks of intraluminal LECs for Static and Flow conditions were obtained from confocal *z*-projections stained for F-actin (phalloidin, green) and nuclei (DAPI, blue) to determine cellular bounding box dimensions (red rectangles). The white arrow depict direction of IF. D) IF induces a reduction in intraluminal cell bounding box height, compared to Static conditions. Dashed lines of violin plot represents medians, and dotted lines denote upper and lower quartiles (*N* > 1000 cells for **–**Flow conditions, and *N* > 300 cells for +Flow conditions). Ad hoc *t*-tests were performed in A) and one-way ANOVA was performed with Tukey pairwise comparisons in B) and D), where * indicates *p*-value < 0.05, ** < 0.01, *** < 0.001, and **** < 0.0001. Data are expressed as mean ± SD. E) Proposed flow-mediated morphological relationship whereby intraluminal cell bounding box height is proportional to vessel diameter.

Control conditions were defined as acellular devices in which cylindrical lumens were not seeded with HDLECs for either Static or Flow conditions (**Figure 6A**). Acellular Static devices did not produce any appreciable change in lumen diameter over the course of 96 hours (∼ 0%). Comparatively, lumens in the acellular Flow devices constricted an average of 7% to 279 μm, primarily occurring within the first 24 hours of implementing IF. Thus, flow-driving IFP is responsible for a modest amount of luminal constriction—although not statistically significant— due to the pressure gradient within the ECM chamber and, potentially, hydrogel swelling effects from sustained IF. While cells independently caused constriction in Static vessels—reaching a stable diameter of ∼ 250 μm by Day 2—additional constriction occurred for vessels experiencing IFP gradients and IF. Flow-induced vessel constriction cannot be solely attributed to the hydrogel effects observed in the acellular Flow control. Thus, there is vasoconstriction by LECs in response to IF and cells are necessary for structural remodeling. The additional constriction under Flow may also be due to the selective lymphangiogenesis occurring at the blind end and the morphogenic interplay between extraluminal sprouting and intraluminal cells. Moreover, we confirmed that there were no significant changes to total intraluminal cell count across the four vessel conditions (**Figure 6B**).

These changes in lumen diameters prompted us to investigate morphological changes of intraluminal cells. After segmenting individual LECs with Cellpose,^[34]^ we used ImageJ to circumscribe a bounding box around each cell to estimate its width and height (**Figure 6C**). With the application of IF, intraluminal cells had a significant reduction in bounding box height compared to Static LECs (**Figure 6D**). Static and Static+VEGF-C intraluminal cells had average bounding box heights of 37 ± 10 μm and 36 ± 11 μm, respectively. In contrast, Flow and Flow+VEGF-C cells had average bounding box heights of 26 ± 8 μm and 29 ± 12 μm, respectively. Thus, IF caused intraluminal cells to reduce their height by approximately 30%— comparing Static and Flow cases. With VEGF-C, IF reduced cellular bounding box height by approximately 19%—comparing Static+VEGF-C to Flow+VEGF-C. In the absence of flow, VEGF-C did not affect cell height, however, in the presence of flow, VEGF-C induced a ∼10% increase in intraluminal cell bounding box height, which is consistent with reports that show VEGF-C to loosen LEC junctions.^[14]^ Taken together, we propose that the flow-induced reduction of intraluminal LEC height is associated with vessel constriction (**Figure 6E**). Namely, the differential arc length of the vessel can be described as r d0, where r is the radius of the vessel and d0 is the differential angle. In this sense, bounding box height (H) is directly proportional to vessel diameter. Given the same number of intraluminal cells, as r d0 reduces, the diameter must also reduce. Therefore, we suggest that morphological changes of intraluminal cells are linked to vessel-level responses mediated by IF.

### 2.7. Intraluminal directionality and elongation of LECs in response to flow

Endothelial cell morphology was further explored by evaluating the directionality of intraluminal LECs. As before, confocal images of F-actin-and DAPI-stained microvessels were utilized to generate *z*-projections, however, we divided LECs into two cohorts based on the Top and Bottom surfaces of lymphatic microvessels (**Figure 7A**). Binary masks of cells were constructed from the *z*-projections to segment individual cells. The directionality of each cell, 0_luminal_, was defined as the angle of the major axis compared to the longitudinal axis of the vessel, which coincided with the direction of intraluminal flow (**Figure 7B**). Static and Static+VEGF-C cells did not have global alignment resulting in roughly uniform distributions and negligible mean resultant vectors. Interestingly, Flow and Flow+VEGF-C cells had a bimodal distribution of alignment with peaks corresponding to either the Top or Bottom surface (**Figure 7C**). Cells under the Flow condition had mean angles of 14° and -13° for Top and Bottom surfaces, respectively, while cells under the Flow+VEGF-C condition had mean angles of 27° and -21° for Top and Bottom surfaces, respectively. Therefore, IF induced the remodeling of intraluminal cells to orient along the vessel axis in the direction of luminal flow, with Flow cells more aligned than their Flow+VEGF-C counterparts. Moreover, to reconcile the bimodality of Top and Bottom cohorts, we propose that intraluminal LECs remodel into a helical structure in the presence of flow (**Figure 7D**). This morphogenic response accounts for how LECs organized in 3-D while aligning in the direction of flow. Interestingly, we consistently observed the Top cohort to have positive angles and Bottom cohort to have negative angles, suggesting there is a driving force that leads to the reliable left handedness of these lymphatic vessels.

**Figure 7.**
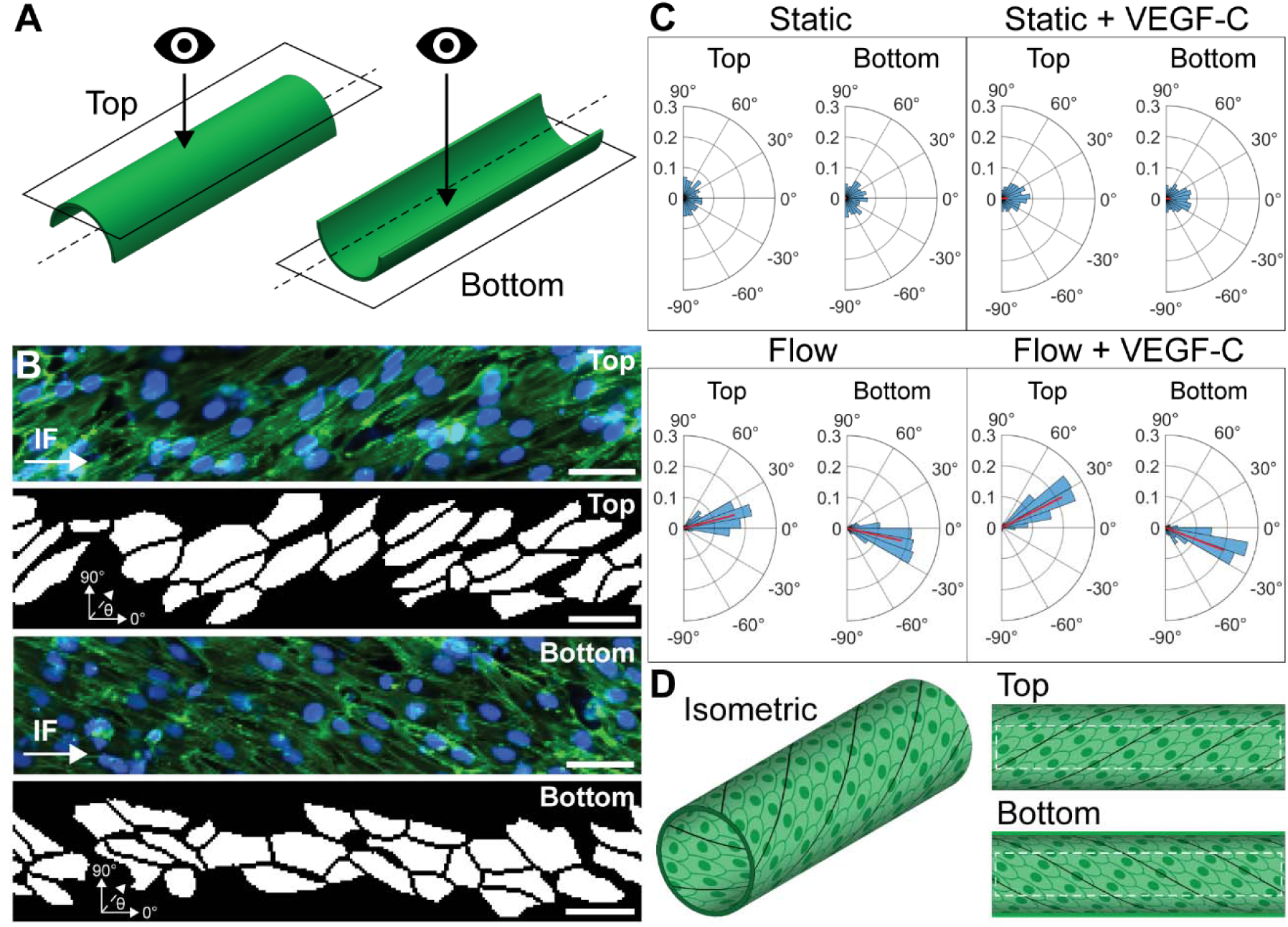
Flow-induced cellular alignment with helical morphogenic response. A) Schematic of Top and Bottom vessel surfaces as obtained by confocal microscopy such that Top is viewed from the outside the vessel, while Bottom is viewed from inside the vessel. B) Top and Bottom confocal *z*-projections of a representative Flow vessel stained for F-actin (phalloidin, green) and nuclei (DAPI, blue), and corresponding cell segmentation. For each cell, 0_luminal_, is defined as the angular difference between the cellular major axis and the longitudinal vessel axis. White arrows depict the direction of intraluminal flow, parallel to the vessel axis. Scale bars are 50 μm. C) Probability-normalized polar histograms of intraluminal cell directionality for the four experimental conditions segmented into Top and Bottom cohorts. Flow induces cellular alignment compared to Static cells. Red lines denote mean resultant vectors (*N* > 1000 cells for **–**Flow conditions, and *N* > 300 cells for +Flow conditions). D) Proposed flow-induced orientation of LECs in a helical pattern to reconcile biphasic Top and Bottom alignment. Dashed white boxes correspond to cropped confocal images in B to reduce effects from 2-D projections. Black lines underscore putative helical morphology.

Given the geometric effect of Top and Bottom surfaces by confocal microscopy (**Figure 7A**), we mirrored Bottom images to harmonize intraluminal cell alignment as viewed from outside the vessel. Upon mirroring, both Top and Bottom cohorts could then be pooled into one population per condition and assessed globally (**Figure 8A**). While Static and Static+VEGF-C cells did not have a strongly coordinated alignment, Flow and Flow+VEGF-C cells had mean angles of 14° and 24°, respectively. Flow and Flow+VEGF-C histograms were also significantly different than a circular uniform distribution—unlike static conditions—further suggesting flow-induced directionality. Additionally, intraluminal cell alignment was analyzed based on the spatial distribution within the vessel (**Figure S6**, Supporting Information). Cells were divided into four subsets based on their distance from the blind end, corresponding to Zones 1–4 of Figure 4. For Flow conditions, mean angles were 18°, 14°, 13°, and 11° for Zones 1–4, respectively. In Flow+VEGF-C conditions, mean angles were 23°, 24°, 25°, and 23° for Zones 1–4, respectively. Thus, in the absence of VEGF-C, LECs become more axially aligned downstream from the blind end. This was expected as luminal flow velocity and shear stress increase from Zone 1 to Zone 4 as the lymphatic vessel collects more interstitial fluid (**Figure S2B**, Supporting Information). Cells appeared less sensitive to variations in luminal velocity and shear stress when VEGF-C was present. Enhanced lateral sprouting due to VEGF-C may account for the differential alignment, reenforcing an interplay between extravascular and intraluminal regions of the vessel.

**Figure 8.**
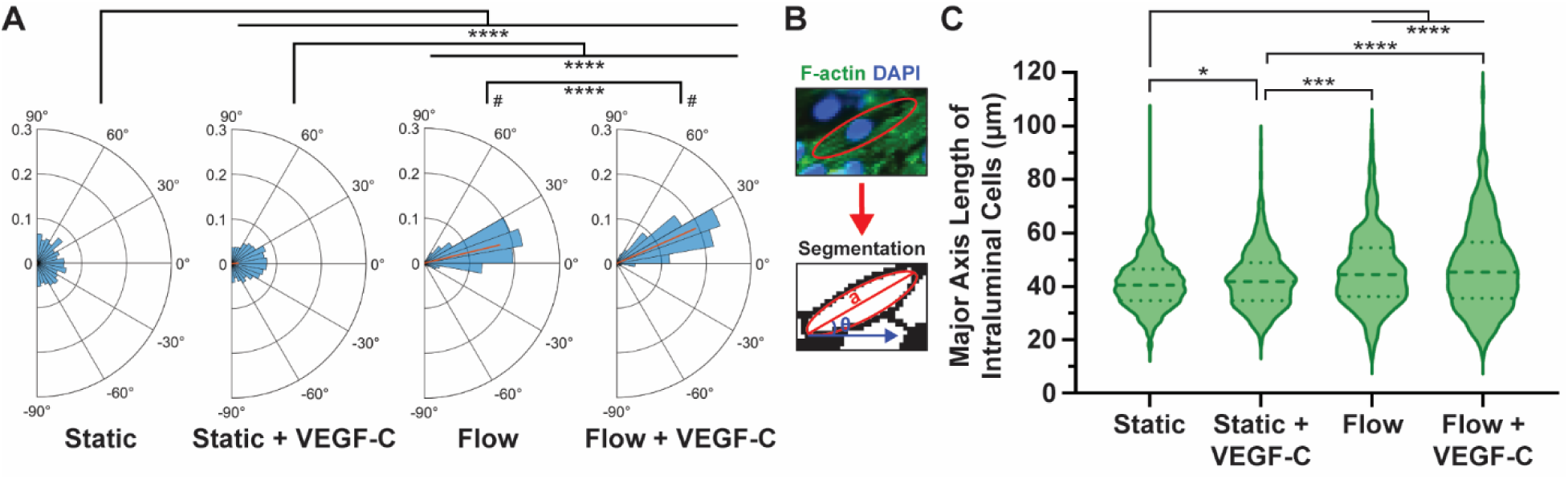
Intraluminal alignment and elongation of LECs in response to flow. A) Probability-normalized polar histograms of intraluminal cell directionality from Figure 7, where Top and Bottom cohorts were pooled, defining directionality as viewed from outside the vessel. Red lines represent mean resultant vectors. Two-sample Kuiper test was performed to compare experimental conditions, and Rao’s spacing test was employed for circular uniformity, where # indicates a *p*-value < 0.05. B) Representative confocal image of intraluminal cells stained for F-actin (phalloidin, green) and nuclei (DAPI, blue), followed by cell segmentation to obtain major axis length, *a*, and major axis angle, 0_luminal_, relative to the longitudinal vessel axis. C) Major axis length of intraluminal cells for the four experimental conditions. Dashed lines in violin plot indicate medians, and dotted lines depict upper and lower quartiles (*N* > 1000 cells for **–**Flow conditions, and *N* > 300 cells for +Flow conditions). One-way ANOVA was performed with Tukey pairwise comparisons, where * indicates *p*-value < 0.05, ** < 0.01, *** < 0.001, and **** < 0.0001.

Lastly, the morphology of intraluminal LECs was evaluated by the major axis length (*a*) of individual cells to identify flow-potentiated elongation (**Figure 8B-C**). An analysis of intraluminal cell major axis length revealed Static and Static+VEGF-C cells to have lengths of 41.1 ± 9.2 µm and 42.8 ± 11.1 µm, respectively. In contrast, Flow and Flow+VEGF-C cells had lengths of 46.1 ± 14.0 µm and 47.4 ± 16.1 µm, respectively. Thus, intraluminal LECs experiencing flow were prompted to significantly elongate, approximately 11%, compared to static conditions.

## 3. Discussion

The most basic function of lymphatics is fluid drainage. Therefore, it is not surprising that forces exerted by interstitial fluid would elicit substantial control over the form and function of lymphatic vessels. Yet, despite tremendous advances in lymphatic-on-chip platforms,^[10b, 36]^ the role of fluid forces in coordinating lymphatic expansion and remodeling is not well understood. By virtue of engineering blind-ended lymphatics and mimicking their biophysical microenvironment, we recreated lymphangiogenesis and evaluated structural remodeling in response to interstitial, transmural, and luminal flow. We believe the present study is the first to recapitulate these flow dynamics for initial lymphatics *in vitro*. Interstitial flow guided sprouting cells within the ECM, transmural flow potentiated lymphangiogenesis, and luminal flow aligned capillary cells in a concurrent, orchestrated fashion. Collectively, our novel MPS integrated intraluminal and extravascular regions of initial lymphatic microanatomy allowing for a coordinated tissue-level response within a 3-D microenvironment.

The pressure gradient within our microdevice drove IF and generated a continuum of transmural flow velocities along the same tubular vessel structure. Given selective sprouting at the blind end, we propose that lymphangiogenesis is strongly correlated with transverse flow velocity across the endothelium of individual microvessels. Our experiments showed that IF on its own was sufficient to mediate a robust lymphangiogenic response. These results align with previous *in vivo* studies where lymphangiogenesis in adult tissue was observed to require IF, and VEGF-C had a reduced ability to rescue lymphatic function in tissue with poor IF.^[37]^ Furthermore, we observed that VEGF-C enhanced overall sprouting when administered with IF, which is consistent with previously reported synergistic effects^[20b]^ and known regulation by VEGF-C.^[35]^ However, the spatial distribution of sprouts along the vessel remained the same in the presence and absence of VEGF-C, suggesting that mechanical flow is the central regulatory cue of lymphangiogenesis while VEGF-C is a stimulant.

In addition to the inductive role of IF on sprouting morphogenesis, IF also oriented nascent sprout growth through the ECM—both in the presence and absence of VEGF-C. Multicellular sprouts grew collinearly to streamlines, against the direction of flow, corroborating previous findings that IF provides directional guidance cues.^[12b, 20b]^ However, in contrast to uniform, one-dimensional IF from literature, sprouting LECs in our model grew collinearly to streamlines within a non-uniform 3-D flow field. Coordinated lymphangiogenesis is particularly relevant within the TME where high pressure gradients at the tumor margin putatively generate elevated levels of IF.^[4]^ Models have shown that IF directs tumor cell migration within the interstitium,^[20a]^ which can potentiate tumoral intravasation into lymphatic capillaries and metastatic dissemination to tumor-draining lymph nodes. Although lymphangiogenesis in solid tumors is generally correlated with metastasis and poor clinical outcomes, a recent study demonstrated the potential of lymphangiogenic induction to enhance immunotherapy and boost T cell immunity through a novel cancer vaccine.^[38]^ In this mouse melanoma model, lymphatic growth occurred locally to irradiated tumor cells overexpressing VEGF-C and promoted *in situ* priming of T cells and lymph nodes for a robust immune response. Therefore, it is possible to harness lymphatic growth to promote tumor-specific immunity. Separately, elevated transmural flow was shown to upregulate programmed death-ligand 1 (PD-L1) expression in tumor-associated blood vessels suggesting a biophysical mechanism in which leaky tumoral vasculatures enable cancer cells to escape immune surveillance.^[39]^ PD-L1 was also shown to regulate immune responses and transendothelial migration of T cells across LECs in 2-D transwell assays.^[39]^ Our lymphatic MPS could serve to model these events in controlled 3-D environments and reveal the differential mechanisms that underlie lymphatic metastasis and antitumor immunity.

With respect to intravascular remodeling, IF was shown to be a morphoregulator and drive endothelial organization into capillary-like lumens over two decades ago using transwell culture models.^[40]^ More recently, microfluidic technologies and organ-on-a-chip devices^[41]^ have made it possible to model vasculature with high fidelity and further unravel the role of fluid flow in 3-D physiological systems. *In vivo*, the collection of IF by initial lymphatics results in the generation of luminal flow or perfusion within the capillary as lymph. Perfusion, in turn, potentiates the alignment and elongation of endothelial cells that comprise the vessel monolayer. Intraluminal LEC alignment has been reported between 28–45°, relative to the vessel axis, for *in vitro* lymphatic vessels^[42]^ and lymphangion^[43]^ subjected to shear stresses between 0.1 and 1.8 dyn/cm^2^. Two important distinctions, however, are that these models were constituted by open-ended vessels with diameters ranging from 400–800 μm, compared to our ~ 250-μm blind-ended vessels. Nevertheless, we present concordant results where static LECs did not align, while intraluminal LECs under Flow and Flow+VEGF-C conditions axially aligned at 14° and 24°, respectively. We hypothesize that enhanced sprouting by VEGF-C accounts for the varying degree of LEC alignment due to the interconnectivity between cells in the extravascular and intraluminal regions. As LECs sprout and invade the ECM, they may pull intraluminal cells with tensile force affecting the alignment angle.

The orientation and elongation of LECs with luminal flow was surprising given the relatively low average shear stress in our vessels: 0.004 dyn/cm^2^, ∼ 25-fold less than the lower end of reported open-ended lymphatic systems.^[43]^ However, luminal flow velocities and accompanying shear stresses in open-ended systems do not recapitulate the flow dynamics of capillary lymphatics; rather, they are representative of downstream collecting lymphatics with muscular contractions.^[2b, 14]^ Notwithstanding the absence of downstream collecting lymphatics, our microvessels retained their drainage function due to the applied pressure gradient. Notably, our MPS drained to zero gauge pressure compared to *in vivo* capillary lymphatics that operate as vacuums due to negative pressures supplied by lymphangions.^[44]^ Moreover, unlike open-ended lymphatic systems, intraluminal flow velocity in our MPS was not uniform: it was lowest towards the blind end and increased moving downstream along the vessel axis to a maximum of ~ 80 μm/s (**Figure S2B**, Supporting Information). Therefore, we believe our microfluidic device is the first to study vascular remodeling and lymphangiogenesis as a result of flow dynamics in capillary lymphatics. Additionally, we note that *in vivo* human lymphatic capillaries are approximately 30–80 μm in diameter,^[3]^ or a fraction of the size of our 250-μm-diameter vessels, and thus physiological shear stress would be significantly higher at the same flow rate.^[27]^ However, templating lumens below 100 μm is an engineering challenge.^[45]^ Alternative approaches to fabricate sub-100-micron lumens include multi-photon lithography^[46]^ and sacrificial microfibers.^[47]^ Despite this limitation of our study, we hypothesize that fluid absorption at the blind end is critical to intercellular alignment at low shear stress.

As a corollary to cellular elongation, bounding box height reduced for intraluminal cells under luminal flow. Compared to Static conditions, IF caused cells to elongate 11% and reduce their bounding box height by 30%. We demonstrate that cellular remodeling due to IF is coupled with a 15% constriction in vessel diameter. We propose that this outcome can be explained, in part, due to the physical geometry of intraluminal cells, such that cellular bounding box height is proportional to vessel radius (H <X r d0). However, we acknowledge that there are also IFP-induced physical effects to hydrogels that contribute to vessel constriction, evidenced by the acellular lumens under IF. Our lymphatic MPS has the capacity to parse the relative contributions of hydrogel swelling and LEC remodeling on IF-mediated vasoconstriction in future iterative studies. For example, in the absence of gel ports, the ECM chamber could be pressurized to varying degrees while maintaining the same net pressure gradient between inlet and outlet ports. This would serve to evaluate IFP-dependent vessel remodeling in the form of constriction or dilation. An alternative explanation for IF-induced constriction is that selective sprouting at the blind end may be pulling the vessel longitudinally: invading tip cells exert tensile force on neighboring stalk cells,^[48]^ which pull intraluminal cells parallel to the vessel axis. In this sense, enhanced sprouting in the Flow+VEGF-C condition may generate additional radial stress on the vessel as lateral sprouts pull orthogonal to the vessel axis, leading to larger diameters than Flow-only treatment. Taken together, we suggest that IF prompts morphological changes of intraluminal cells, which is linked to vascular constriction and extraluminal sprouting.

In addition to axial alignment and elongation, flow dynamics coordinated intraluminal LECs to organize into a helical structure. A consistent left-handed helical pattern was observed for IF-induced lymphatic microvessels, defined by the direction of luminal flow. The helical organization of recapitulated blood vessels has been noted,^[49]^ but is largely overlooked for 3-D tissues. However, in recent years with the aid of microfabrication technology, cell chirality was proposed as an intrinsic property of the cell, mediated by helical actin filaments within the cytoskeleton.^[50]^ Consequently, actin-driven cell chirality may account for tissue asymmetry^[51]^ and the helicity observed in our lymphatic microvessels. In a separate study, cell chirality was exploited to affect the barrier function of human umbilical vein endothelial cells (HUVECs). The loss or randomization of cell chirality led to the disruption of intercellular junctions and the dysregulation of endothelial permeability.^[50b]^ While these findings suggest that cell chirality is an important mediator of endothelial function, the experiments were conducted on 2-D micropatterned substrates and endothelial permeability was measured by 2-D transwell assays. The findings and methods described in the present study have the potential to be widely utilized to assess endothelial cell chirality and helical morphogenesis boosting our mechanistic understanding of these cellular-and tissue-level phenomena. Given our results, we hypothesize that flow dynamics are required for LEC chirality to manifest at the vessel level in blind-ended lymphatics. Furthermore, disorganized and loosely connected endothelial cells are characteristic of leaky tumor vessels.^[52]^ Thus, by affecting LEC chirality, it would be possible to recapitulate discordant tumor-like lymphatic microvessels in our system.

## 4. Conclusion

We engineered blind-ended lymphatic microvessels and reconstructed their biophysical microenvironment to assess cellular and vascular remodeling. By incorporating physiologically relevant IF and controlling the hydraulic resistance of the ECM, we mimicked functional lymphatic capillaries and induced lymphangiogenesis over the course of four days. Selective sprouting at the blind end was strongly correlated with transverse flow velocity across the endothelium of individual microvessels. Extravascular sprouting through the ECM was guided by IF, while intraluminal cells responded with coordinated alignment and elongation. We observed morphological changes at the cellular level that propagated to tissue-level helicity and vasoconstriction. Furthermore, intraluminal morphogenesis was coupled to extraluminal sprouting due to flow dynamics and subsequent fluid collection. We conclude that there is a substantial interrelationship between intra-and extraluminal regions of blind-ended lymphatic vessels due to local flow-induced structural remodeling. By recapitulating capillary blind-end lymphatic structure and fluid absorption function *in vitro*, our MPS is positioned as a powerful tool to advance our understanding of lymphatic biology and pathophysiology that will contribute to the development of therapies that target lymphangiogenesis.

## 5. Experimental Section

### 5.1 Cell culture

Commercially available HDLECs (PromoCell, C-12217) were purchased and maintained using EGM-2 MV cell culture media (Lonza, CC-3202). Cell passages of 5–10 were used in this study. Cells were cultured in T-75 flasks (Greiner Bio-One, 658175) within a humidified incubator at 37°C and 5% CO_2_ with media exchange every two days. Cells were harvested from flasks by washing with 1× Dulbecco’s phosphate-buffered saline (PBS) without Mg^2+^ and Ca^2+^ (Gibco, 14190250) followed by detachment using 0.05% Trypsin-EDTA (Gibco, 25300054) for 3–4 minutes. After neutralizing trypsin with 10% FBS in DMEM, HDLECs were centrifuged at 220 RCF for 3 minutes and resuspended in cell media at a concentration of 10 x 10^3^ cells per μL in preparation for seeding the lumens of the microfluidic devices.

### 5.2 Fabrication of the microfluidic devices

Microvessel devices were fabricated using xurography-based techniques established by our lab^[25]^ (**Figure S1**, Supporting Information). Briefly, a single device was comprised of five layers of PDMS that were irreversibly bonded to each other by plasma oxidation (Harrick Plasma, PDC-32G). Individual layers were formed by spin coating (Specialty Coating Systems, 6812P) PDMS on silanized silicon wafers (University Wafer, 100 mm) at a 10:1 ratio of base to curing agent (Dow, 184 Silicone Elastomer Kit). Layers were spun at 170 rpm and 115 rpm to generate thicknesses of 250 µm and 400 µm, respectively. Spin-coated PDMS layers were then cured overnight in a 65°C oven. The 400-µm-thick layer was subjected to xurography (Graphtec, CE7000) to create a 750-µm-wide rectangular channel extending from the inlet to the back of the device; thus, a ∼ 250-µm-diameter nitinol wire (Malin, 0.01-inch diameter) could be inserted into the microchannel. During the layer-by-layer assembly, biopsy punches (Integra) were used to bore circular ports (e.g., inlet, outlet, and gel ports) and the central hydrogel chamber. The 6-mm-thick top-most layer of the device was formed by pouring PDMS pre-polymer into a 100-mm Petri dish to the specified height. A relatively thicker top layer permitted sufficiently deep inlet and outlet ports to hold adequate volumes of media for robust culture, minimizing osmolality effects due to evaporation. Additionally, a thicker device facilitated the press-fit attachment of a liquid hopper for Flow devices. After sealing the fifth PDMS layer to enclose the gel chamber, the assembled device was then bonded to a 150-µm-thick glass slide (VWR, 48393-106) for future microscopy. The completed assembly was stored in a 100-mm deep-dish culture plate (Fisherbrand, FB0875711) and sterilized with UV light for 30 minutes before subsequent steps.

For Flow devices, the upstream channel length was cut to 8 mm by xurography, as opposed to 1 mm for Static devices to provide additional hydraulic resistance and control IF. Furthermore, a liquid hopper was constructed from a sterile 20-mL syringe (BD, 302830): the plunger was discarded, and the barrel of the syringe was hand-sawed to create an open-top liquid reservoir. Furthermore, the exterior threading of the Luer lock of the syringe was cut to expose the interior tip of the syringe. The diameter of the tip of the liquid hopper permitted a friction-based press fit connection with the 4-mm-diameter inlet port of the microdevice. Cell culture media was then pipetted into the liquid hopper to the specified height to generate a 20 mmH_2_O pressure head and induce IF. Lastly, we implemented a liquid ramp at the outlet to usher media out of the device allowing it to pool at atmospheric pressure on the glass surface.

### 5.3 Formation of blind-ended lymphatic vessels

To enhance gel-PDMS attachment,^[53]^ the device was pre-coated with 1 mg mL^-1^ of polydopamine (PDA) for one hour at room temperature (RT) and rinsed three times with 1× PBS. PDA solution was prepared by mixing dopamine hydrochloride (Sigma-Aldrich, H8502) with 10 mM, pH 8.5 tris-HCl buffer (Bioworld, 420204141) and subsequent sterile filtration (Thermo Scientific, 0.22 μm, PES). PDA treatment was critical to facilitate the durable anchorage of collagen to the PDMS surfaces of the microdevice, thus ensuring proper IF through the gel. Following the PDA coating, the nitinol wire was inserted into the microchannel of the device such that the end of the wire stopped at the midplane of the 4-mm-diameter central gel chamber; the wire served to form a hollow cylindrical blind-ended lumen structure upon casting collagen gel (**Figure 1A-B**). Type I collagen (Corning, 354249) isolated from rat tail was prepared to a working concentration of 3 mg mL^-1^ at a pH of 7.4 from 10× PBS, 1 N sodium hydroxide (Sigma-Aldrich, S2770100ML), and sterile cell-grade water (VWR, 02-0200-0500), per manufacturer’s instructions. Gels were pre-incubated at 4°C for approximately 12 minutes before casting to enhance fiber formation.^[54]^ Subsequently, collagen was pipetted into the central ECM compartment of the microdevice via the 1.5-mm-diameter gel ports and allowed to polymerize for at least 30 minutes at 37°C in a humidified incubator. After polymerization, the wire was removed by vacuum aspiration to template a cylindrical lumen within the gel chamber for endothelial cell seeding. The back channel—from which the wire was removed from the device—was then plugged with PDMS to prevent leakage at the outlet port, and the device was stored in the humidified incubator.

Prior to cell seeding, the cylindrical lumens were coated in 100 μg mL^-1^ human fibronectin (Sigma-Aldrich, FC010) for 30 minutes to improve cellular attachment and biocompatibility of the device. After flushing with media, 2 μL of HDLEC-laden media, suspended at a concentration of 10 x 10^3^ cells per μL, was pipetted into the blind-ended lumens at the base of the outlet port. To improve cell adhesion, devices were first placed upside down in the incubator for 15 minutes within hydrated Petri dishes before inverting. Lumens were then flushed with culture media to clear excess unattached cells. HDLECs were cultured within the microdevice one day prior to the implementation of IF or treatment with VEGF-C to stabilize the vessels without affecting initial cell attachment. For growth factor conditions, recombinant human VEGF-C (PeproTech, 100-20CD) was added to cell media at a concentration of 50 ng mL^-1^. Lymphatic microvessels were imaged daily by phase contrast microscopy (Invitrogen, EVOS XL Core) to monitor and evaluate day-to-day changes (**Figure S3B**, Supporting Information).

### 5.4 Immunofluorescence

Blind-ended lymphatic microvessels were stained within our MPS by modifying existing immunofluorescence techniques from our lab.^[55]^ Briefly, at Day 4, microvessels were fixed with 4% paraformaldehyde (PFA) for 30 minutes at RT, permeabilized with 0.2% Triton X-100 for 30 minutes at RT, blocked with blocking buffer for 30 minutes at RT, and subsequently stained for VE-cadherin (monoclonal antibody, BD Biosciences, 561567), F-actin (phalloidin, Fisher Scientific, A12379), and nuclei (DAPI, Sigma-Aldrich, D9542). Microvessels were washed three times between each step with washing buffer: 0.1% Tween-20 in 1× PBS. Blocking Buffer consisted of 1% BSA in washing buffer. VE-cadherin was stained overnight at 4°C, F-actin was stained for 60 minutes at RT, and DAPI was counterstained for 10 minutes. Immunofluorescence images were acquired on a Nikon AXR confocal microscope controlled with NIS-Elements AR Software. 3-D renders from confocal *z*-stacks were generated for each of the four experimental conditions (**Videos 1-4,** Supporting Information).

### 5.5 Quantification of sprouting area

Upon immunofluorescent staining and confocal microscopy, blind-ended lymphatic microvessels were visualized by 2-D maximum intensity *z*-projections. Images of F-actin-and nuclei-stained vessels were cropped to confine the vessel length to 1000 μm for standardization. Images were then binarized and a mask of each lumen was generated using analysis software, ImageJ. The lumen mask was subtracted from the binarized image to isolate lymphatic sprouts in white on a black background. Sprouting area was measured by ImageJ’s Analyze Particles plugin. For each vessel, the average vessel diameter was determined from the area of the lumen mask. Sprouting area was normalized to the surface area of each vessel to account for vessels of varying diameters.

Sprouting area was further analyzed by zonal distribution. A theoretical blinded-ended vessel was taken to be a 250-μm-diameter cylinder with a closed front face and an open back face. Therefore, the *z*-projection of a theoretical vessel was a rectangle with a width of 250 μm and length of 1000 μm. Given the theoretical diameter, zones were partitioned with commensurate widths of 250 μm and taken to be normal to the vessel surface. Zone 0 was defined at the area normal to the front face of the blind-ended lumen, and Zones 1–4 were defined at the areas normal to the vessel surface along the longitudinal axis. Due to the asymmetry of the blind-end, Zone 0 and Zone 1 were separated by 45° angles taken to be normal to the corners of the rectangular lumen such that a 90° angle was formed within the vessel at its midline (**Figure 5A**). Zonal sprouting area was normalized to the corresponding vessel surface area within each zone.

### 5.6 COMSOL multiphysics model and analysis

The 3-D geometry of the microvessel and device were constructed in COMSOL Multiphysics 6.2 with precise dimensions. The model was comprised of three distinct material domains: cell media, collagen, and lymphatic cells. The Laminar Flow interface was used to define the physics and solve for velocity and pressure of fluid within the device in addition to wall shear stress along the vessel. An inlet boundary condition was set on the top surface of the inlet port of the device with a constant pressure and fully developed flow. An outlet boundary condition was set on the top surface of the outlet port of the device set to zero gauge pressure, fully developed flow, and prevent backflow. The top surfaces of the two gel ports were treated as zero flux boundaries, which was supported by the observation of no fluid flow out of these ports during experiments.

To optimize the design of the Flow device, a parametric sweep of the upstream channel length was performed along with an auxiliary sweep of the inlet pressure (**Figure 2C**). The values used for upstream channel length were 0.5, 1, 2, 4, 8, and 16 mm. The values used for inlet pressure were 0, 10, 20, 30 mmH_2_O. The cell media material properties were defined using the built-in properties of water. Since this study was done prior to the calibration of collagen and cell domain properties, values from literature were used. The collagen material was defined by a porosity of 0.94 and a permeability of 0.41 µm^2^, the latter being an average of reported values.^[21]^ HDLECs were similarly defined with custom values for porosity and permeability. To ensure that the lymphatic cellular domain was a function of permeability and not porosity, the porosity value was set to 1. A cellular permeability of 0.01 µm^2^ was used based on the average reported LEC hydraulic conductivity^[21]^ of 1×10^-5^ cm^3^/dyn·s, assuming a solvent viscosity of 1 mP and cell layer thickness of 10 µm. The steady state solution was then calculated for every parameter combination. Transverse flow velocity was extracted by taking the surface average of the normal velocity component (***u***) on the interior surface of the vessel. Ultimately, an 8 mm upstream channel length tolerated a wider variation of inlet pressure to achieve 3 μm/s compared to shorter channel lengths, which required a narrower range of pressure for the same velocity. Therefore, an 8 mm channel effectively led to a larger acceptable margin of error in inlet pressure to achieve the desired velocity; this was preferential given the drop in pressure within the liquid hopper over 24 hours, albeit relatively small: < 1 mmH_2_O. Thus, the microdevice with an 8 mm upstream channel and a 20 mmH_2_O pressure head was employed throughout this study as our Flow device.

To calibrate the collagen and cell parameters of the model, experimental pass-through volumes of media were compared to analytical FEA results for three different configurations of the Flow device: no lumen (lumenless), a lumen without HDLEC cells (acellular), and an endothelialized lumen (vessel). Collagen permeability was calibrated from flow rates in the lumenless and acellular configurations (**Figure 2E**). Steady state solutions for the two models were computed with an auxiliary study extension for collagen permeabilities ranging from 0.2 to 0.4 µm^2^. The total pass-through volume was extracted by integrating the normal velocity component of the fluid over the outlet surface. Once average collagen permeability was determined, this value (0.304 μm^2^) was used in the vessel model for subsequent FEA studies. Since the experimental flow rate for the vessel was found to be approximately the same as the acellular lumen, a finite permeability for the LECs could not be determined. Instead, the upper bound found in literature of ∼0.1 µm^2^ was selected. This permeability was then used for further vessel models.

IF streamline values were extracted from the FEA results at the centroid location of each sprouting cell and tabulated in a .txt file. Centroid (x, y) coordinates, previously determined from Cellpose and ImageJ analysis, were used to specify probe points within the FEA model for data export. The steady state solution of the Flow device with the calibrated material parameters was used along with a vessel diameter of 250 µm. Since experimental vessels varied in diameter, the individual cell centroids were adjusted in the y-direction, multiplying by the ratio of theoretical to experimental vessel diameters. Streamline angles were then compared to the major axis of sprouting LECs to determine the relative orientation of cells within the collagen matrix (**Figure 5**).

### 5.7 Empirical flow rate

For Flow devices, the volumetric flow rate was measured to validate the predicted results from COMSOL analysis. Flow rate was determined by taking the average pass-through volume of media over 24 hours. Liquid hoppers were filled with media to a height of 20 mm above the top of the microdevice, and liquid ramps were initiated with 100 μL of media. Pass-through volume was calculated by averaging two methods: i) pooled media at the outlet was collected and weighed, and ii) the quantity of media required to refill the liquid hopper to its initial height was measured. The averaging of these two methods was performed to minimize bias due to evaporation as each method represents the minimum and maximum possible pass-through volume, respectively.

### 5.8 Cell segmentation using Cellpose

Cellpose, an open-source cellular segmentation algorithm,^[34]^ was employed to segment two categories of cells: i) intraluminal cells within the lymphatic microvessel, and ii) extraluminal sprouting cells that had invaded the collagen-based ECM. For intraluminal cells, two *z*-projections were made for each vessel from F-actin-and nuclei-stained confocal *z*-stacks, corresponding to the top and bottom of the vessel surface (**Figure 7B**). To avoid boundary effects at the vessel edges due to the 2-D projections, images were cropped to rectangles centered at the midline of the vessel. For extraluminal cells, a single *z*-projection was created for the entire vessel, and a rectangular lumen mask was removed to isolate sprouting cells. These images were imported into the Cellpose graphical user interface (GUI). Subsequently, the built-in cyto3 model was run with F-actin and DAPI as channels 1 and 2, respectively, to generate masks of individual cells. Auto-generated masks were manually reviewed, and additional masks were drawn to append any cells that the cyto3 model had not identified. For each vessel, the collection of cell masks was saved as a NumPy file for further analysis in Python. Custom Python scripts were written to output binary cellular mask data after applying two filters: cells that touched the boundary of the binary images were excluded as they were considered incomplete masks, and cells below an area threshold of 78 μm^2^ (i.e., rr(5 µm)^2^) were omitted to avoid including fragmented cells. Binary cellular mask images were then opened with ImageJ, and the Analyze Particles plugin was used to calculate and tabulate individual cell morphological data: centroid location, area, major axis angle, major axis length, and bounding box height. The invasion depth of extraluminal sprouting cells was quantified by measuring the shortest distance between the cell centroid location and the rectangular lumen mask. The major axis angle of extraluminal cells was used to quantify its relative angle with respect to FEA streamlines.

### 5.9 Sprouting alignment to interstitial flow streamlines

Theoretical IF streamline vectors were calculated from COMSOL analysis based on the microdevice and blind-ended geometry, individual vessel diameters, experimentally measured collagen permeability, and the hydraulic conductivity of the HDLECs^[21]^. Streamline angles were extracted from the FEA results for each coordinate within the collagen gel corresponding to the centroid location of sprouting cells, as determined from Cellpose and ImageJ. Major axis angles of cells were presented using MATLAB’s quiver plot mapped to their respective location on the *x-y* plane. Subsequently, the difference between the streamline angle and the major axis angle was calculated for each invading cell. Polar histograms were normalized by probability with 10-degree-wide buckets ranging from -90 to 90 degrees, where 0° represents the streamline. For sprouting cells under the Static+VEGF-C condition, cell orientation was taken with respect to projected streamlines to be able to directly compare to IF conditions. Additionally, the morphology of invading LECs in the Static+VEGF-C condition was assessed by comparing the rounded phenotype (i.e., aspect ratio < 1.5) to cells under IF conditions (**Figure S5**, Supporting Information).

### 5.10 Statistical Analysis

Numerical data reported in this manuscript are expressed as mean ± standard deviation (SD). Each experimental condition was performed with at least four replicate microvessels. Statistical analysis was conducted by either student’s *t*-test or analysis of variance (ANOVA) with pairwise Tukey comparisons using GraphPad Prism 10. The details of data representation are provided in the corresponding figure captions. Circular statistics were applied to polar directionality data by the CircStat toolbox^[56]^ in MATLAB: Rao’s spacing test was used for circular uniformity and the Kuiper test was employed to determine if two samples differed significantly. The following notation was used to compare statistical difference: ∗ for *p*-value < 0.05, ∗∗ < 0.01, ∗∗∗ < 0.001, and ∗∗∗∗ < 0.0001; # for *p*-value < 0.05 for circular uniformity.

## Competing Financial Interests

Jonathan W. Song is a co-founder of and shareholder in EMBioSys, Inc.

## 6. Acknowledgements

The authors acknowledge support from an NSF CAREER award (CBET-1752106), the Mark Foundation for Cancer Research (18-024-ASP), the National Heart Lung Blood Institute (R01HL141941), and the Ohio State University (OSU) Materials Research Seed Grant Program, funded by the Center for Emergent Materials, an NSF-MRSEC, grant DMR-1420451, the Center for Exploration of Novel Complex Materials, and the Institute for Materials Research. One of the authors, J.C.H., gratefully acknowledges funding from the OSU Pelotonia Graduate Fellowship program and support from an NHLBI Graduate Diversity Supplement. S.S.A. is funded through the OSU Fellowship and the Ohio State Distinguished University Fellowship. J.W.T. and J.M.B. gratefully acknowledge support from the OSU Pelotonia Undergraduate Scholars Program. Confocal microscopic images presented in this report were generated using instruments and services at the Campus Microscopy and Imaging Facility (CMIF) at OSU. This facility is supported in part by grant P30 CA016058, National Cancer Institute.

