## Supplemental Figures and Captions for "Fluid forces control structural remodeling of blind-ended lymphatic microvessels"

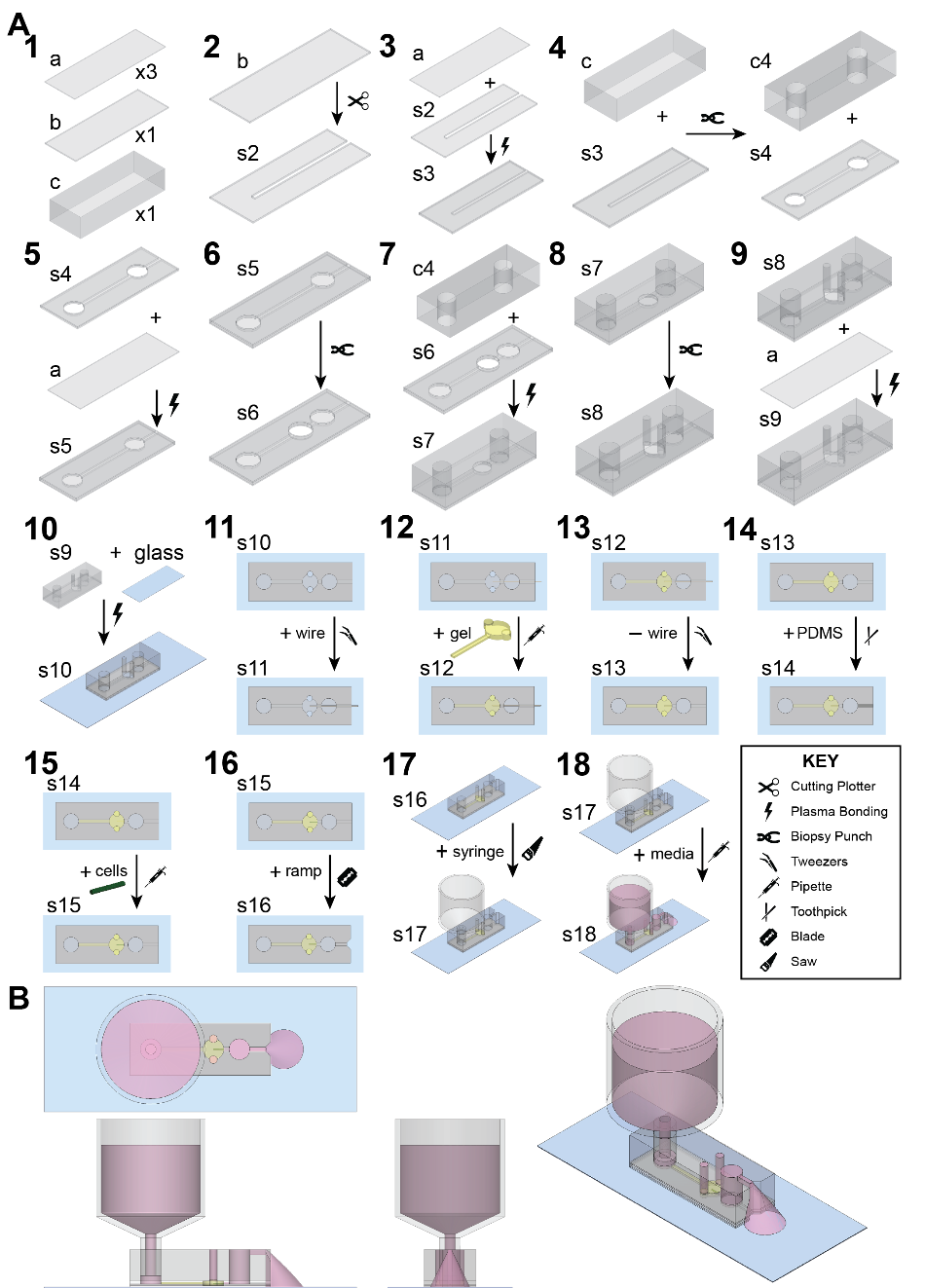

**Supplemental Figure 1.** Microfabrication and assembly of the novel microphysiological system to create blind-ended lymphatic microvessels. A) Stepwise fabrication of the PDMS-based microdevices by benchtop xurography. Depicted in Step 1, the device is created from five total layers: (i) three 250-µm-thick layers (part a), (ii) one 400-µm-thick layer (part b), and (iii) one 6-mm-thick top layer (part c). The length of the microchannel (part s2) cut in Step 2 by a cutting plotter determines whether the assembly will be for a Static or Flow device: Static and Flow devices have 1- and 8-mm upstream channel lengths, respectively. The layer-by-layer assembly is conducted with irreversible plasma bonding. The central ECM chamber, inlet, outlet, and gel ports are made from biopsy punches. Upon templating a cylindrical lumen that terminates within a tunable hydrogel, lymphatic endothelial cells are seeded to recapitulate a blind-ended microvessel within a biophysical ECM. For Flow devices, an inverted syringe is press fitted into the inlet port and filled with cell culture media to generate a controllable hydrostatic pressure head and drive interstitial flow. A ramp is cut at the outlet to allow perfused media to pool out the back of the device, analogous to electrical grounding. B) Orthographic projections of a fully constructed Flow microdevice bonded to a glass slide: top, side, front and isometric views (counterclockwise from upper left).

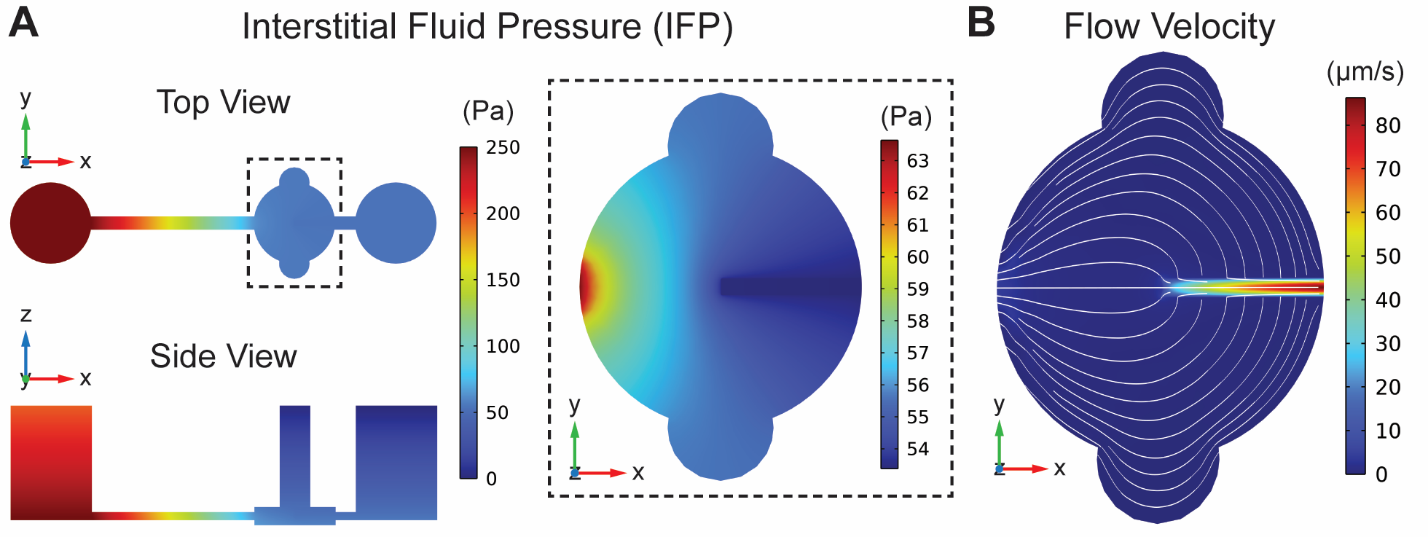

**Supplemental Figure 2.** Interstitial fluid pressure (IFP) and flow velocity within the Flow microdevice predicted by finite element analysis. A) Top View at the mid *z*-plane of the vessel and Side View of fluid pressure within the microdevice. IFP is greatest at the inlet due to the hydrostatic pressure head specified by the liquid hopper. The 8-mm-long hydrogel-filled upstream microchannel provides the principal resistance to flow in the microdevice; as such, the drop in IFP mostly occurs within the upstream channel. Inset highlights the moderate pressure gradient within the ECM chamber that generates physiological interstitial flow (IF) velocity on the order of 1–5 μm/s (see Figure 5). B) Top view of fluid flow velocity within the ECM chamber and lymphatic microvessel at the mid *z*-plane of the vessel. White lines depict left-to-right IF streamlines. Within the vessel, intraluminal velocity is lowest at the blind end—at magnitudes similar to IF—and accelerates downstream as fluid uptake accumulates in the less resistive luminal space.

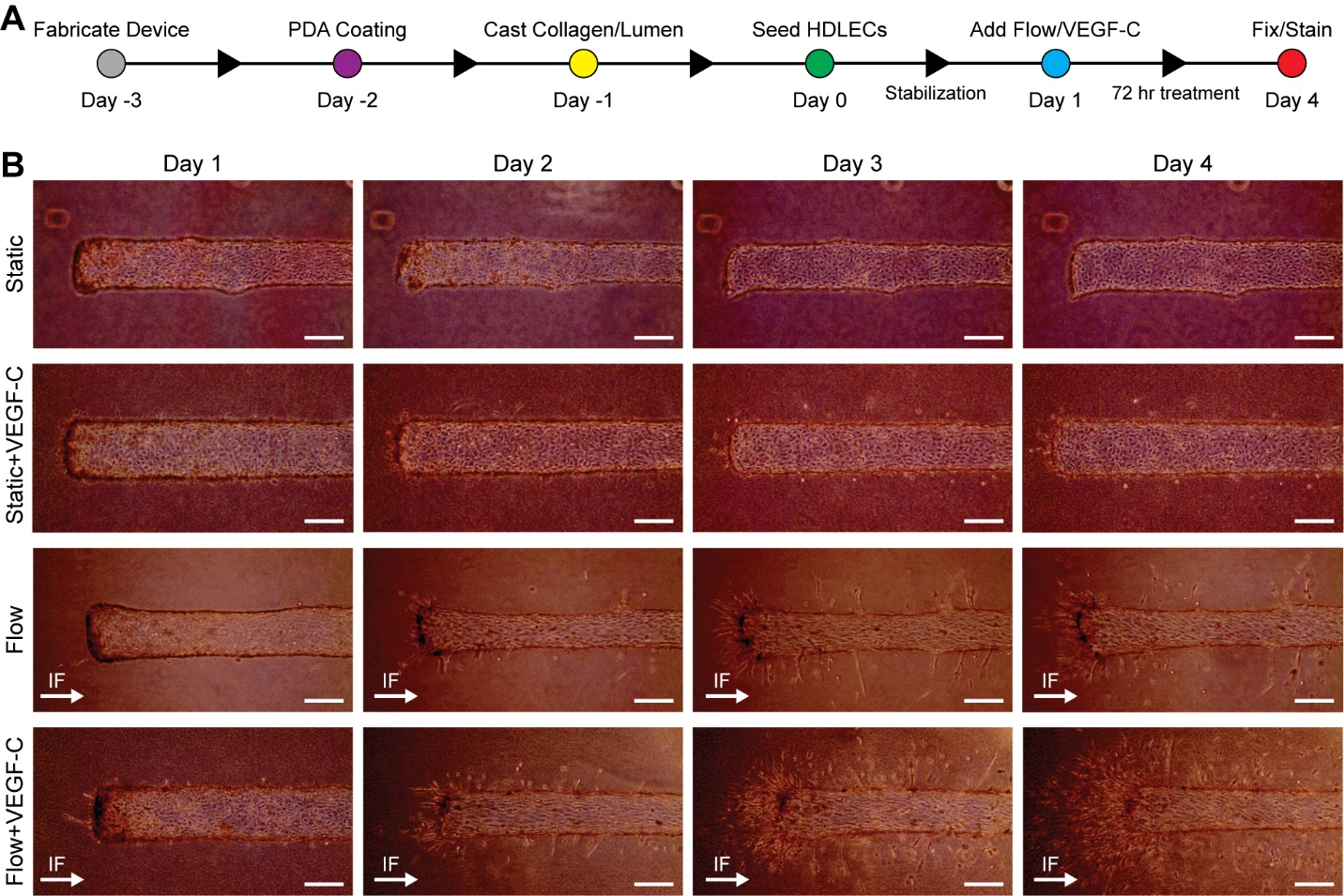

**Supplemental Figure 3.** Daily progression of lymphatic microvessels. A) Experimental workflow consisting of device preparation, HDLEC seeding at Day 0, administration of interstitial flow (IF) and/or VEGF-C at Day 1, and 72 hours of subsequent treatment. B) Representative phase-contrast images demonstrating the day-to-day progression of lymphangiogenesis and morphological changes by blind-ended lymphatic vessels subjected to the four experimental conditions within 3 mg mL^-1^ collagen gel. White arrows indicate the direction of IF. Scale bars are 200 µm.

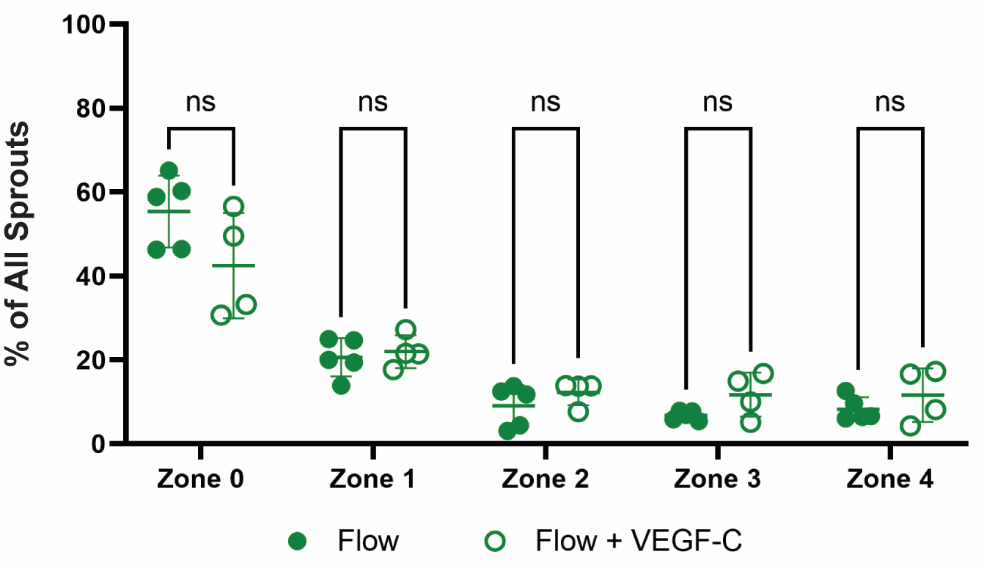

**Supplemental Figure 4.** Zonal distribution of sprouting morphogenesis as a percentage of all sprouting activity. VEGF-C augments sprouting overall but does not affect the spatial distribution by zone. Sprouting area was normalized as a percentage of respective vessel surface area within each zone and then divided by total normalized sprouting area across all zones. Sprouting per zone is presented for Flow conditions with and without VEGF-C (*N* = 4 or 5 vessels). Data are expressed as mean ± SD. Two-way ANOVA was performed with Tukey comparisons where “ns” indicates non-significance, *p*-value > 0.05.

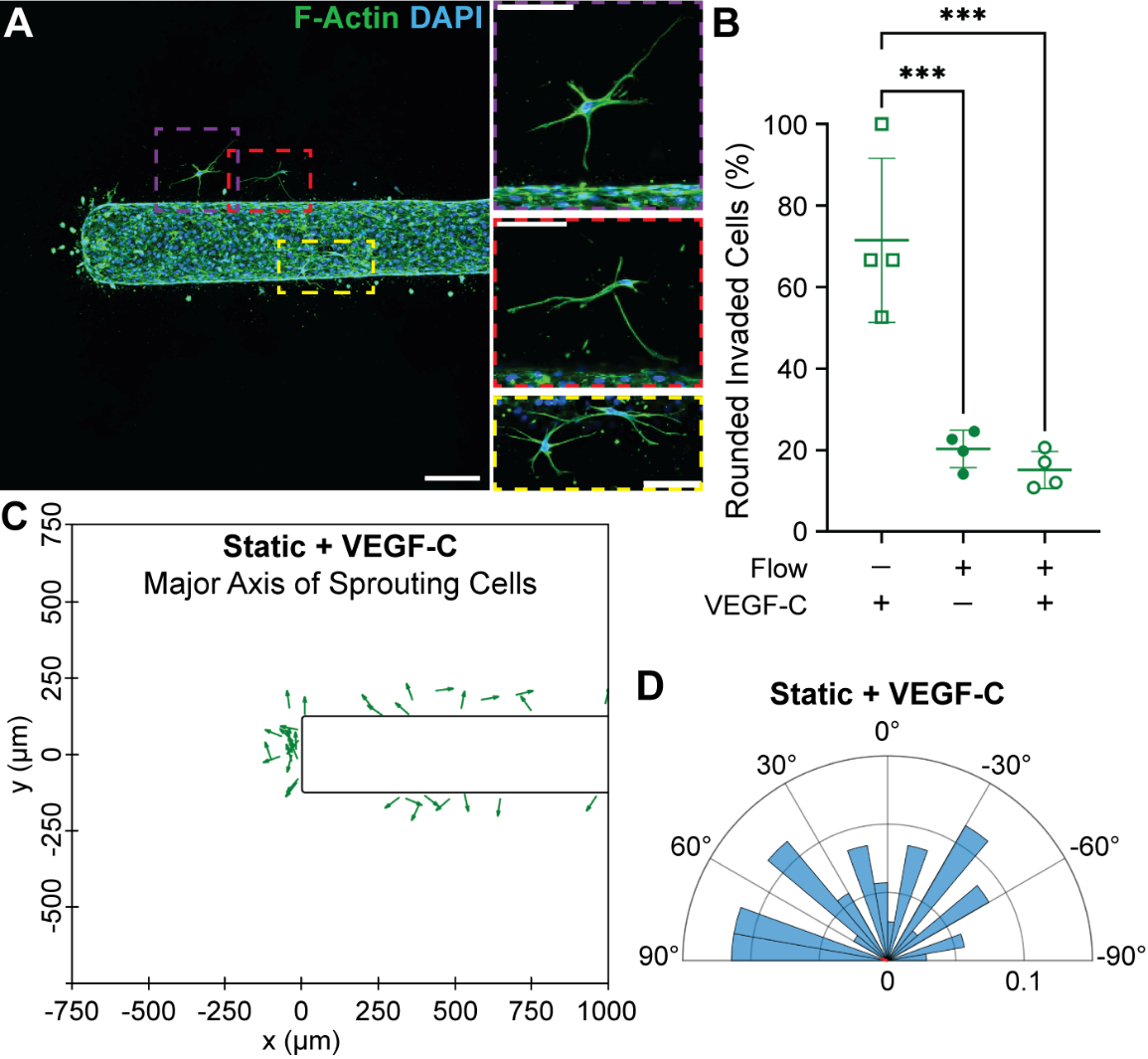

**Supplemental Figure 5.** Morphology of invading LECs under the Static+VEGF-C condition. A) Confocal *z*-projection of representative lymphatic vessel stained for F-actin (phalloidin, green) and nuclei (DAPI, blue), fixed at Day 4. Scale bar is 200 μm. Three insets depict a subset of sprouting cells with an unusual “star-shaped” morphology in the Static+VEGF-C condition characterized by multiple, long, actin-rich extensions stretching in various directions. Inset scale bars are 100 μm. B) Percentage of sprouting cells per vessel with a rounded phenotype, defined by an aspect ratio < 1.5. A majority of invading cells in the Static+VEGF-C case were rounded compared to +Flow conditions, suggesting that interstitial flow (IF) is needed to sustain sprouting and connectivity with the main vessel. C) The major axes of individual sprouting cells are plotted as green arrows and mapped to their respective locations within the collagen-based ECM (*N* = 4 vessels). D) Probability-normalized polar histogram of the major axes of spouting cells from panel C (*N* > 30 cells). Cell orientation was taken with respect to projected streamlines to directly compare with IF conditions. The small red line depicts a near-zero mean resultant vector implying random directionality. Thus, in the absence of IF, sprouting LECs have no coordinated orientation.

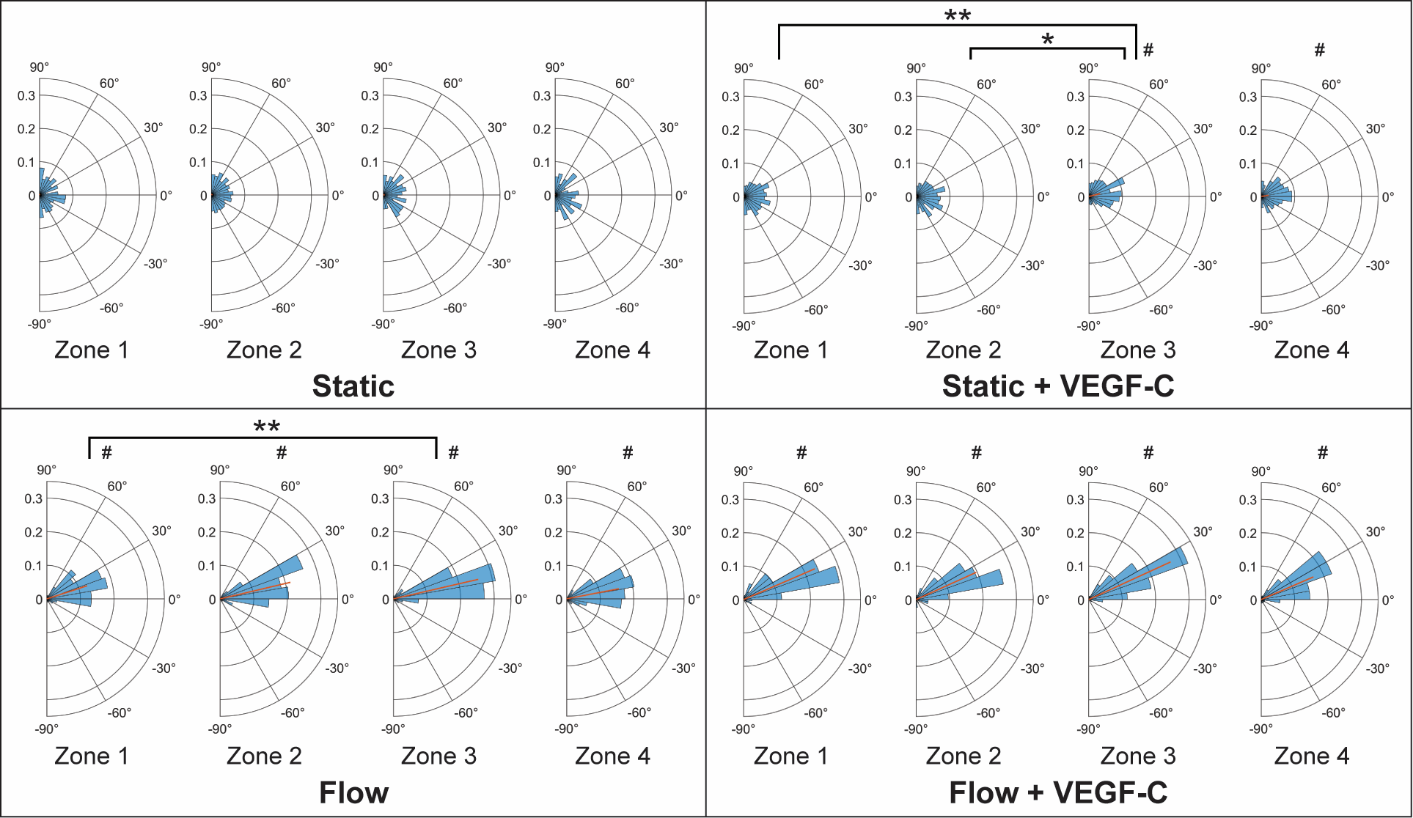

**Supplemental Figure 6.** Intraluminal zone-wise alignment of LECs. For the four experimental conditions, the angle of the major axis of cells is reported per zone with respect to the vessel axis (0°), as defined in Figure 8. Luminal cell populations were spatially subdivided by their distance from the blind end, i.e., by Zones 1–4 defined in Figure 4. Red lines depict mean resultant vectors (*N* > 1000 cells for **–**Flow conditions, and *N* > 300 cells for +Flow conditions). Static and Static+VEGF-C cells did not have coordinated alignment to the vessel axis. Zonal analysis revealed that intraluminal Flow cells became more aligned downstream from the blind end, consistent with increased luminal velocity and wall shear stress: mean angles of 18° and 11° for Zones 1 and 4, respectively. Flow+VEGF-C cells appeared less sensitive to variations in luminal velocity and shear stress. Two-sample Kuiper test was performed to compare zones, where * indicates a *p*-value < 0.05 and ** < 0.01. Rao's spacing test was employed for circular uniformity, where # indicates a *p*-value < 0.05.
